# Multi-omics causal deep learning reveals regulatory structure while interventions refine perturbation predictions

**DOI:** 10.64898/2026.11.22.768440

**Authors:** Shervin Goudarzi, Vikram Nandi, Nuo Liu, Hani Goodarzi

**Affiliations:** Department of Electrical Engineering and Computer Sciences, University of California, Berkeley, Berkeley, CA, USA; College of Computing, Data Science, and Society, University of California, Berkeley, Berkeley, CA, USA; Arc Institute, Palo Alto, CA, USA; Department of Biochemistry and Biophysics, University of California, San Francisco, San Francisco, CA, USA

**Author notes:** **Corresponding authors** Correspondence to: Nuo Liu and Hani Goodarzi.

## Abstract

Inferring gene regulation from single-cell data is difficult because observational measurements cannot distinguish correlation from causation and access to perturbational data, especially with multiple modalities, remains limited. Here we present the Cross-Modal Causal Variational Autoencoder (CMC-VAE), which learns a network linking genes and proteins (gene–gene, protein–protein and gene–protein) in the latent space and maps each connection back onto measured features. Trained on 809,130 observational CITE-seq profiles, CMC-VAE recovers co-expression, transcription factor regulons and immune signaling programs, and learns a sparse cross-modal map enriched for transcript–protein cognate pairs. On three CRISPR screens used during development, it ranks knockout-responsive genes better than correlation and latent-VAE baselines. Fitted to a screen’s unperturbed control cells, without perturbation labels, the model translates measured knockout RNA responses into protein responses better than a correlation map and cognate transcripts. The direction of gene-to-gene regulation, however, is not recovered from observational data. On the CITE-seq screens, fine-tuning only the latent network improves held-out response correlation, mainly by learning a response shared across knockouts. On larger Perturb-seq screens, with about 1,000 training perturbations, fine-tuning the latent network learns gene-specific effects that transfer to held-out genes, more so than a gene-level causal model fine-tuned on the same cells, and in simulation, such interventions also restore edge orientations that observational training leaves unresolved.

## Introduction

Cells respond to infection, signalling cues and genetic perturbations through coordinated changes in RNA and protein abundance. Understanding these responses requires knowing which genes influence other genes and how those effects reach proteins. CITE-seq is a multimodal technique that measures RNA and surface proteins in the same cell, providing a way to study these relationships together.^1^ Yet RNA and protein levels can vary together due to one regulating the other, both responding to a shared signal, or marking the same cell type. Paired measurements therefore reveal relationships without necessarily establishing which feature acts upstream. Understanding and disentangling these causal relationships beyond pure covariation in biology remains an unsolved problem.

Genetic perturbations help distinguish these possibilities by testing what changes when a gene is disrupted. However, perturbation screens cover only a subset of genes and cellular conditions, and screens that measure paired RNA and protein information are rare. This raises a practical question: how much useful structure can a model learn from abundant observational measurements, and how much additional perturbation data is needed to improve its predictions? Causal inference theory explains why some observational models cannot distinguish one direction from another, while stronger assumptions can make direction recoverable in particular settings through observational data.^2–6^ Whether those assumptions hold for single-cell datasets is unclear, so we test several of them here.

Existing methods address different parts of this problem. totalVI, MOFA+ and MultiVI learn representations that integrate modalities and support tasks such as denoising and imputation.^7–9^ HALO studies relationships between chromatin accessibility and transcription using a framework informed by pseudotime.^10^ Methods such as sVAE and NODAGS-Flow use perturbation data to learn causal representations or regulatory relationships.^11,12^ Recent perturbation models also use curated biological knowledge to improve prediction across contexts, as in PIE.^13^ Here, we ask whether a model of paired RNA and surface-protein measurements can learn an explicit network of both translation or feedback (i.e. across modalities) and regulatory (i.e. within modalities) relationships from purely observational data, and then improve its perturbation predictions by updating that network with limited supervision.

We developed CMC-VAE, a latent-based causal VAE architecture that parameterises the graphical structures connecting RNA and protein latent variables through four sets of interactions: within RNA, within protein, RNA to protein and protein to RNA. A linear decoder maps these interactions to scores for measured gene and protein pairs,^14^ while allowing cyclic feedback loops in the latent graph and its projection onto gene- and protein-level graphs. We use “causal” for the latent graph in the sense of a structural parameterisation, but, as we show, its orientation must be supplied by interventions. Trained on an observational corpus of 809,130 paired profiles, CMC-VAE recovered co-expression, known immune programs and a map linking many transcripts to their encoded cognate proteins. We evaluated its predictions in three CRISPR CITE-seq screens and in a fourth screen withheld from model development, using in silico perturbations that edit learned latent graph edges. In the screens used for development, our model ranked differentially expressed genes above several baselines, and this ranking was reduced when the learned latent graph was perturbed. Furthermore, the RNA-to-protein map was enriched for cognate pairs and tracked how well each protein can be read from its transcript, and, fitted to a CRISPR screen’s unperturbed control cells, the model translated knockout RNA responses into protein responses. The observational RNA network itself, however, was nearly symmetric and did not predict the direction of RNA responses. Fine-tuning only the latent graph on a small number of perturbations improved held-out RNA response correlation, which was largely due to learning a response shared across knockouts. With larger Perturb-seq screens, fine-tuning the latent graph learned small gene-specific effects for held-out genes, exceeding a gene-level model fine-tuned on the same cells but not supervised predictors.

## Results

### A latent network connects RNA and protein measurements

CMC-VAE represents each cell using 128 RNA and 64 protein latent variables (Fig. 1a). Separate encoders estimate inputs to a linear structural model, whose adjacency contains connections within each modality and in both cross-modal directions. A linear decoder reconstructs the 2,692 RNA features and 144 surface-protein measurements and maps the latent connections to interaction scores for measured feature pairs (Fig. 1b; Methods). The model permits feedback loops and is trained with a decoder group-lasso penalty and a spectral-stability penalty. The latent graph comprises the within-modality blocks Arr and App and the cross-modal blocks Wrp (RNA to protein) and Wpr (protein to RNA) (Fig. 1b). The latent graph A is represented by equation (1) in Methods. The endogenous latent solves the linear structural equations *z* = *A*^T^*z* + ε, which, rearranged, yields *z* = (*I* − *A*^T^)^−1^ ε. Each block is the product of a differentiable binary gate (Gumbel-sigmoid relaxation) and a real-valued weight. A linear decoder *x̂* = *Dz* + *b* reconstructs both modalities back to the normalized counts so that each latent network can be projected onto the gene or protein space for interpretation (Fig. 1b).

**Fig. 1.**
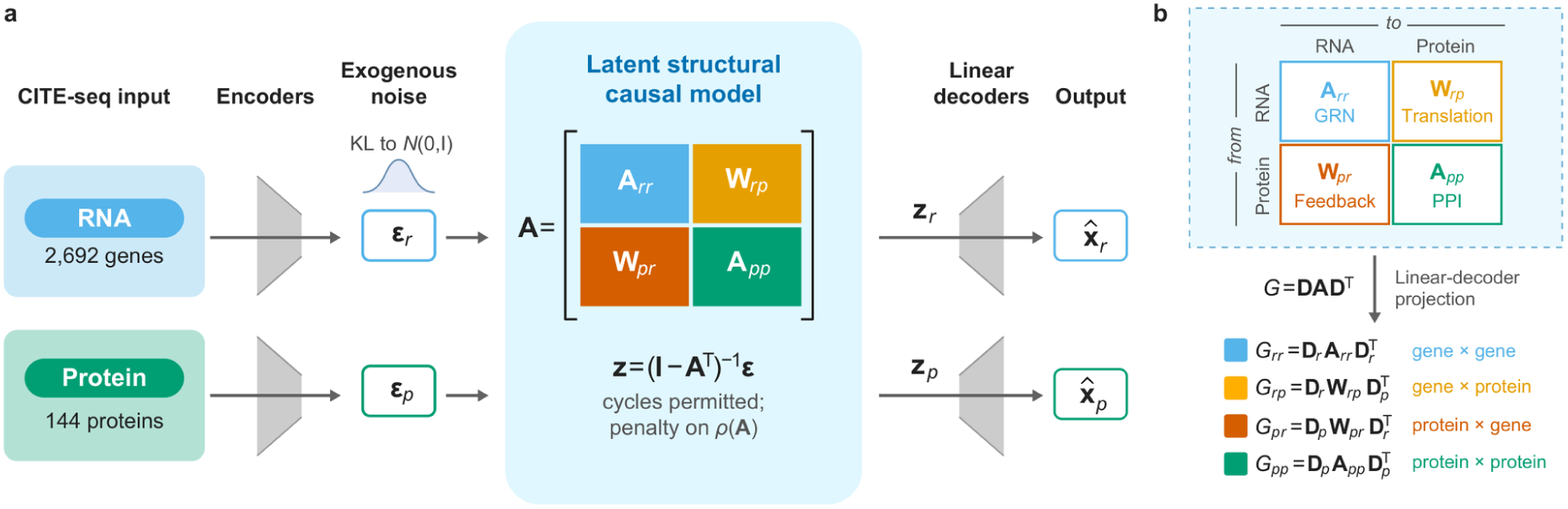
CMC-VAE connects RNA and protein representations through a latent graph. **a**, Separate encoders map paired RNA and surface-protein measurements from peripheral blood mononuclear cells (PBMCs) to distributions over exogenous latent variables. A linear structural model produces endogenous latent variables, which are decoded to reconstruct each modality. The graph permits feedback loops and is trained with a stability penalty. **b**, The adjacency contains the RNA block Arr, protein block App, RNA-to-protein block Wrp and protein-to-RNA block Wpr. The diagram labels the within-modality blocks as a gene regulatory network (GRN) and protein–protein interaction (PPI) network; these are hypothesized roles assessed by the downstream evaluations. Rows index source latents and columns index target latents. The linear decoder maps these blocks to feature-pair scores through G = DADᵀ. The projected scores are evaluated against biological references and perturbation responses.

We fitted the model using a corpus of 809,130 paired peripheral blood mononuclear cell profiles from the Hao and Stephenson CITE-seq studies.^15,16^ The corpus includes a healthy reference and samples from patients with COVID-19. We also compared different training cohorts and cohort combinations. The evaluated settings and selection procedure are reported in Supplementary Note 2 and Supplementary Table 1. Perturbation performance on three CRISPR screens (two Cas9-knockout and one CRISPRi) guided this choice, so these screens also informed model development; a fourth screen was therefore withheld from all development and used only for a final test.

### The learned maps capture known gene relationships and transcript–protein pairs

We first compared the projected RNA network with co-expression and curated reference networks. Its AUROC was 0.98 for co-expression, 0.76 for the STRING reference, 0.68 for DoRothEA and 0.63 for CollecTRI (Fig. 2a).^17–19^ CMC-VAE scored numerically higher than totalVI^7^, GRNBoost2^20^ and DeepSEM^21^ on the two directed regulon references. However, absolute correlation gave similar regulon scores, showing that agreement with these directed databases can largely reflect co-expression. The projected RNA network was also highly sparse as 99% of its edge weights were below 0.1% of the largest weight compared with 98% for GRNBoost2, 75% for DeepSEM and 9% or less for correlation and totalVI (Fig. 2e), a sparsity encouraged by the decoder group-lasso penalty.

**Fig. 2.**
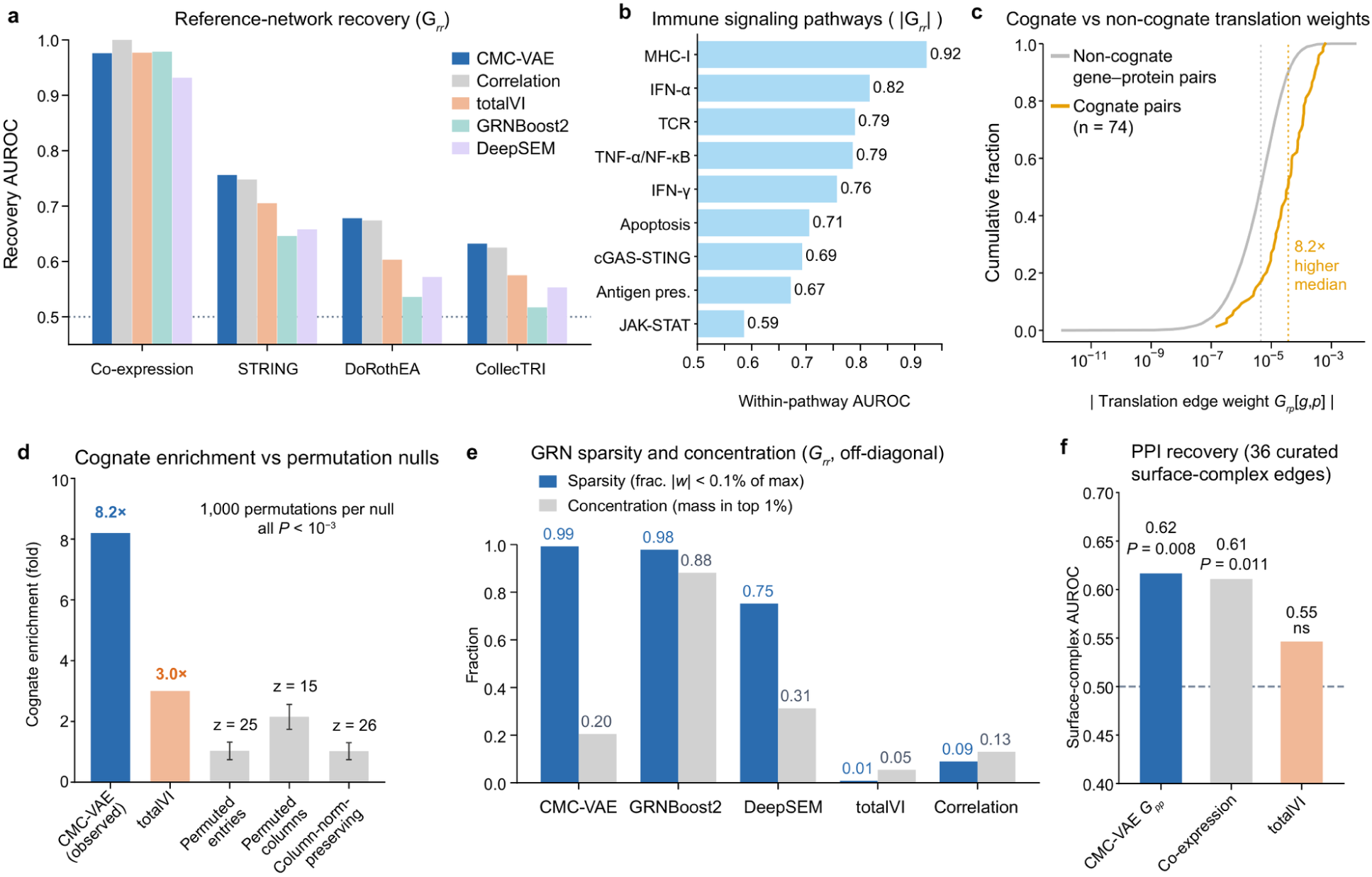
Learned feature scores recover known relationships and cognate RNA–protein pairs. **a**, AUROC against co-expression, STRING and the DoRothEA and CollecTRI regulon references. The co-expression reference is built from correlation, so the correlation baseline reaches an AUROC of 1. **b, Within-program AUROC, ranking projected RNA-network connections among the gene pairs of nine curated immune programs; MHC class I antigen presentation reached an AUROC of 0.92. Panels a and b show the final model (seed 42). c**, Cumulative distributions of absolute RNA-to-protein scores for 74 cognate pairs and the non-cognate background; dotted lines show medians. **d**, Cognate enrichment for CMC-VAE and the totalVI-derived map, with three randomized null distributions for CMC-VAE: permuting entries of the projected RNA-to-protein score matrix, permuting protein-column assignments, and a column-norm-preserving null (which does not preserve graph degree). Null bars show mean ± s.d. over 1,000 randomizations. z denotes the observed statistic’s distance from the null mean in standard-deviation units. **e**, Fraction of RNA-network weights below 0.1% of the largest absolute weight and fraction of total absolute weight in the largest 1% of edges. **f**, AUROC for 36 curated surface-complex connections (15 cis within-complex and 21 trans receptor–ligand pairs across 15 complexes), ranked among all 10,296 protein pairs: CMC-VAE 0.62, marginal co-expression 0.61 and totalVI 0.55; for the 15 cis pairs, CMC-VAE 0.71 and co-expression 0.69. Horizontal reference lines show an AUROC of 0.5. Significance is a one-sided label-permutation test (20,000 permutations): all-edge P = 0.008 (CMC-VAE), 0.011 (co-expression) and 0.18 (totalVI); cis-only P = 0.002 (CMC-VAE) and 0.007 (co-expression). These assess each score against chance, not CMC-VAE against co-expression. Cognate score distributions were compared using a one-sided Mann–Whitney U test (P = 5 × 10⁻¹⁷); because transcript–protein pairs are not independent, the randomized nulls are the primary test. For the randomized nulls, none of 1,000 values reached the observed statistic.

The RNA network ranked connections within nine curated immune programs above chance, including antigen presentation, interferon signalling and T cell receptor signalling (Fig. 2b). The strongest result was for MHC class I antigen presentation, with an AUROC of 0.92. Training separate models on healthy and COVID-19 cells changed the program composition: the model trained on disease cells devoted fewer latent variables to antigen presentation and more to type I and II interferon signalling (Supplementary Note 3 and Supplementary Figs. 1 and 2).

We next examined cognate pairs, in which a measured transcript encodes its corresponding measured protein. Across 74 such pairs, the median absolute RNA-to-protein weight was 8.2 times the median for non-cognate pairs (Fig. 2c). None of 1,000 randomizations under each of three null models produced enrichment as large as the observed value (Fig. 2d; Methods). The corresponding totalVI-derived map showed 3.0-fold enrichment. CMC-VAE’s map was also more concentrated, with a Gini coefficient of 0.74 compared with 0.54 for totalVI (Supplementary Table 2). Thus, the model learned a strong preference for many biologically matched RNA–protein pairs without being given those pairings as edge labels. Raw RNA–protein correlation in the training cells gave a similar enrichment (8.2-fold) but a far less concentrated map (Gini coefficient 0.63; 4% versus 84% of entries below 0.1% of the largest weight; Supplementary Note 15), a concentration that partly reflects the decoder group-lasso penalty. The cognate transcript was, however, never the single top-ranked transcript for its protein (correlation, 12% of proteins; median cognate rank 9.1% versus 3.4% for correlation). CMC-VAE thus learns, from observational training alone, an explicit and sparse map enriched for transcript–protein cognate pairs.

We evaluated the protein–protein map against 36 curated surface-complex connections represented on the antibody panel. It achieved an AUROC of 0.62, similar to protein co-expression at 0.61 and higher numerically than the totalVI-derived score of 0.55 (Fig. 2f). For the 15 direct physical connections, the corresponding values were 0.71 for CMC-VAE and 0.69 for co-expression. These results provide limited support for surface-complex recovery: the effect was not consistent across independently trained models (AUROC 0.46 and 0.57 for the replicate models) and largely reflects co-expression in the data (Supplementary Note 4).

### CMC-VAE ranks knockout-responsive genes in CRISPR screens

We evaluated the observationally trained model in three CRISPR screens: Frangieh, Chen and Arce.^22–24^ For each target gene, a decoder-guided change to the latent graph produced a predicted response, which we compared with the measured difference between perturbed cells and non-targeting control cells from the same condition, estimated by pseudobulk differential expression (Fig. 3a; Methods, “Decoder-guided perturbation prediction”). **AUROC** measured how well predicted effect magnitudes ranked differentially expressed (DE) features. **Signed pearDE** measured the Pearson correlation between predicted and observed changes across those features. Because knockouts in the same screen share a common response component, we also report shared-response references and mean-centred metrics (Supplementary Note 15).

**Fig. 3.**
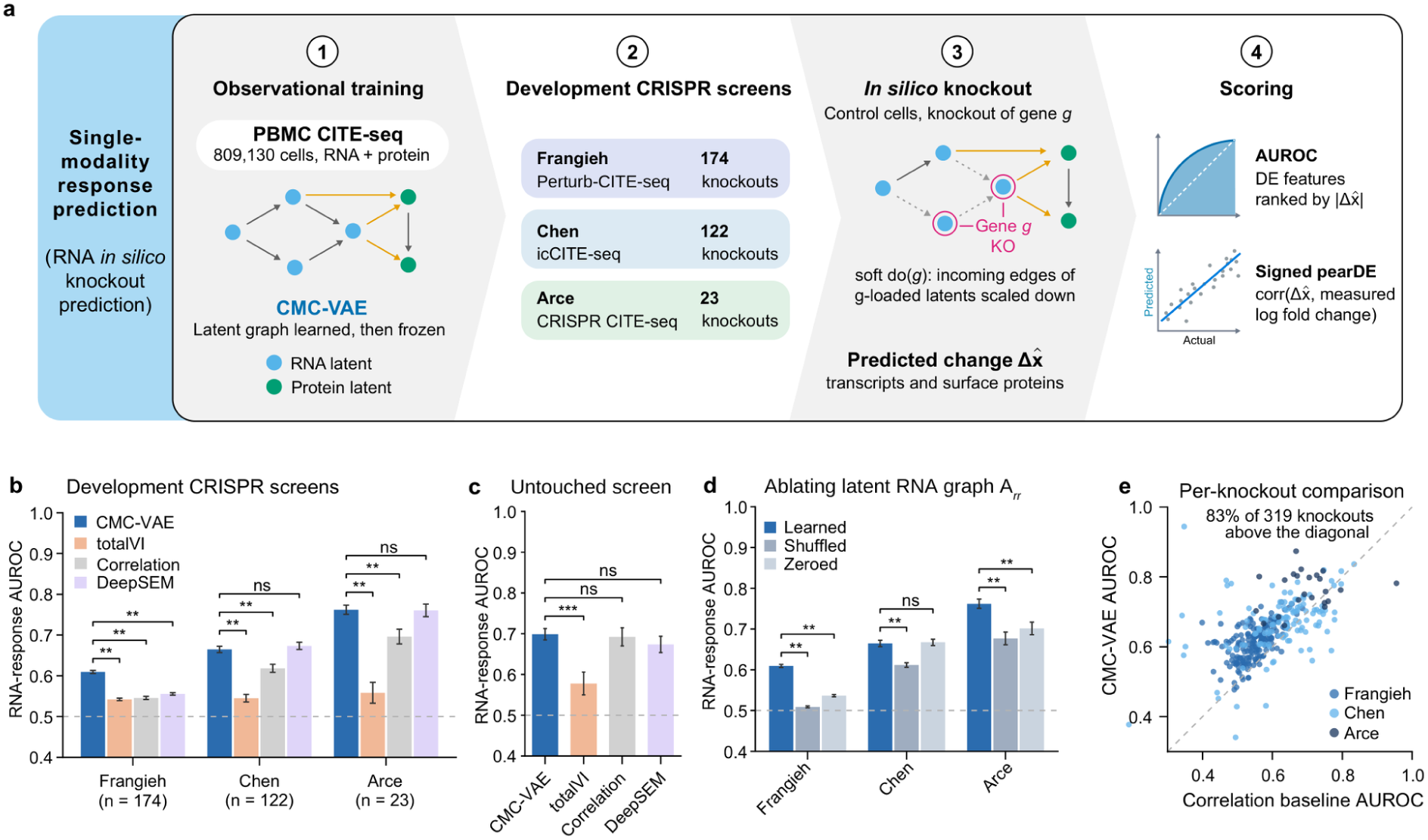
CMC-VAE ranks knockout-responsive genes using its learned latent graph. **a,** Schematic of the perturbation prediction analysis in the three CRISPR CITE-seq screens used during development (Frangieh, Chen and Arce; 174, 122 and 23 evaluated knockouts). For each target gene, a decoder-guided edit of the latent graph (Methods) produces a predicted response, which is compared with the measured response from condition-matched pseudobulk differential expression. **b,** RNA-response area under the receiver operating characteristic curve (AUROC) for ranking differentially expressed (DE) genes, for CMC-VAE, totalVI, the correlation baseline and spectrally stabilized DeepSEM trained on the same cells. **c,** The same analysis in the Papalexi ECCITE-seq screen (THP-1 cells; 13 evaluable knockouts), which was withheld from all model development; values are means over three independently trained models of each method. AUROC was a secondary end point; the pre-specified primary end points (mean-centred pearDE and retrieval) were null for all methods (Supplementary Note 15). **d,** RNA-response AUROC of CMC-VAE with the learned RNA latent graph A_rr_, a shuffled A_rr_ or a zeroed A_rr_, with encoders and decoder unchanged. **e,** Per-knockout AUROC of CMC-VAE against the correlation baseline for the 319 knockouts in b; dashed line, identity; one knockout with a correlation-baseline AUROC of 0.19 lies outside the plotted range. The development screens in b informed model selection (Methods). In b, d and e the final model (seed 42) is shown. Bars show mean ± s.e.m. across knockouts; dashed horizontal lines, AUROC = 0.5. Brackets in b report the more conservative of a joint bootstrap over knockouts and control cells (999 replicates, DE recomputed) and a paired two-sided t-test with Benjamini–Hochberg correction across Fig. 3; brackets in d, joint-bootstrap P values with Benjamini–Hochberg correction within the panel; brackets in c, paired two-sided t-tests with Benjamini–Hochberg correction. Evaluable units are listed in Supplementary Table 7, and signed and protein-response metrics are shown in Supplementary Fig. 8. *P < 0.05, **P < 0.01, ***P < 0.001; ns, not significant.

In these development screens, which informed model selection (Methods), CMC-VAE ranked DE genes above chance for RNA-to-RNA targets in every screen (AUROC = 0.61, 0.66 and 0.76 for Frangieh, Chen and Arce, respectively) and exceeded both the correlation baseline and totalVI in all three screens (Fig. 3b; bootstrap P ≤ 0.004, and P < 0.05 for each of three independently trained models). It was competitive with spectrally stabilized DeepSEM trained on the same data, outperforming it on Frangieh (P = 0.001) and not differing significantly on Chen or Arce. Across the 319 evaluated knockouts, CMC-VAE ranked responsive genes better than the correlation baseline for 83% (Fig. 3e). To test generalization beyond the screens used during development, we applied the final models without any change to the Papalexi ECCITE-seq screen in THP-1 cells,^25^ which had been withheld from all model development. CMC-VAE again ranked responsive genes above chance (AUROC = 0.70; 13 evaluable knockouts), exceeded totalVI (0.58; q = 2 × 10⁻⁴) and matched the correlation baseline (0.69) and DeepSEM (0.67) (Fig. 3c). The pre-specified primary end points on this screen, which measure knockout-specific responses (mean-centred pearDE and retrieval), did not differ from chance for CMC-VAE or for any baseline (Supplementary Note 15).

Signed RNA pearDE remained small and inconsistent in sign for CMC-VAE (0.10, −0.13 and 0.01 in Frangieh, Chen and Arce) and the baselines, so the observational model predicts which genes respond rather than the direction of their change (Supplementary Fig. 8a). Protein-response ranking was also near chance in all three screens, in which the measured surface-protein panels changed little under the screened perturbations (Supplementary Fig. 8b,c and Supplementary Note 15). Rankings by control-cell expression variability, which use no perturbation information, ranked responsive genes even more highly (Supplementary Fig. 9), and predicted responses were more similar across knockouts than measured responses (Supplementary Note 15). Much of the ranking therefore reflects which genes are generally responsive rather than which genes respond to a particular knockout, so we tested the contribution of the learned network directly by perturbing it.

We tested whether these predictions depended on the learned graph by shuffling or removing the connections of the RNA latent block A_rr_ while keeping the encoders and decoder fixed (Fig. 3d). Shuffling A_rr_ reduced RNA-response AUROC in all three screens (by 0.10, 0.05 and 0.09 in Frangieh, Chen and Arce; q = 0.002), and removing it reduced AUROC in Frangieh (by 0.07) and Arce (by 0.06; in two of three independently trained models), with no change in Chen. Transposing the block changed AUROC by at most 0.02 (Supplementary Note 15), consistent with the nearly symmetric RNA graph described below. Because the encoders and decoder were trained together with the graph, these ablations show that the trained model relies on its latent graph for ranking, rather than that the graph alone carries this information; they provide no evidence for a learned orientation of gene–gene relationships.

### Observational training did not recover signed direction for perturbation responses

The perturbation and edge-reversal analyses indicated that RNA-to-RNA direction is not learned from observational training, consistent with classical identifiability theory^2,3,5^. Furthermore, the projected GRN was nearly symmetric as the median forward (TF-to-target) weight matched the reverse weight to within 0.5%, and edge reciprocity was approximately 0.99, compared with approximately 0.01 for a degree-preserving null. This failure was consistent across the evaluated models and training configurations.

We tested whether this result depended on particular modeling choices. Across the 27 configurations reported in Supplementary Table 4, changes to regularization, decoder constraints, nonlinear structural equations, latent dimensions and initialization did not produce strong signed RNA-response prediction in the Frangieh evaluation. Directed priors increased network asymmetry without improving the evaluated response predictions. Additional variants with non-Gaussian priors, independence penalties and auxiliary-variable conditioning also failed to improve RNA-response prediction consistently across cohorts (Supplementary Notes 5–7). Some conditional priors reduced protein-response accuracy. Thus, network asymmetry could be induced, but the direction of held-out RNA responses was still not predicted.

These experiments show the limits of the tested training procedures. They do not establish that direction is impossible to infer from observational data under every model. Classical identifiability results rely on specific assumptions about the graph, disturbances and measurement process.^2–6^ In CMC-VAE, the decoder and latent graph are learned together, adding uncertainty about how a fitted latent connection should be interpreted. We discuss these assumptions in Methods and use the held-out perturbation experiments to assess predictive usefulness directly. Theory for linear causal representation learning indicates that, when the decoder is learned together with the graph, latent edge orientations generally require interventional data,^26–28^ which we test below.

### CMC-VAE learns a cognate translation map and translates knockout responses across modalities

The most specific structure learned from observational training was the cross-modal translation map. This map was cognate-specific (Fig. 2c,d), with its strongest edges connecting canonical lineage markers to their transcripts, including CD19, CD20 (MS4A1), CD22, IgD and IgM (B cells), CD14, CD16, CD36 and CD64 (monocytes) and CD8 (T cells). Cognate edge strength tracked how well each protein was predicted by its own transcript across the 74 cognate proteins (Spearman’s ρ = 0.90; Fig. 4b and Supplementary Note 9). The reverse protein-to-RNA map was similar to the transpose of the RNA-to-protein map (Pearson’s r = 0.71), and after removing the fitted transpose component, the residual rarely ranked receptor-specific target sets better than protein–gene co-expression (Supplementary Note 10), providing little evidence here for independent receptor-specific feedback.

**Fig. 4.**
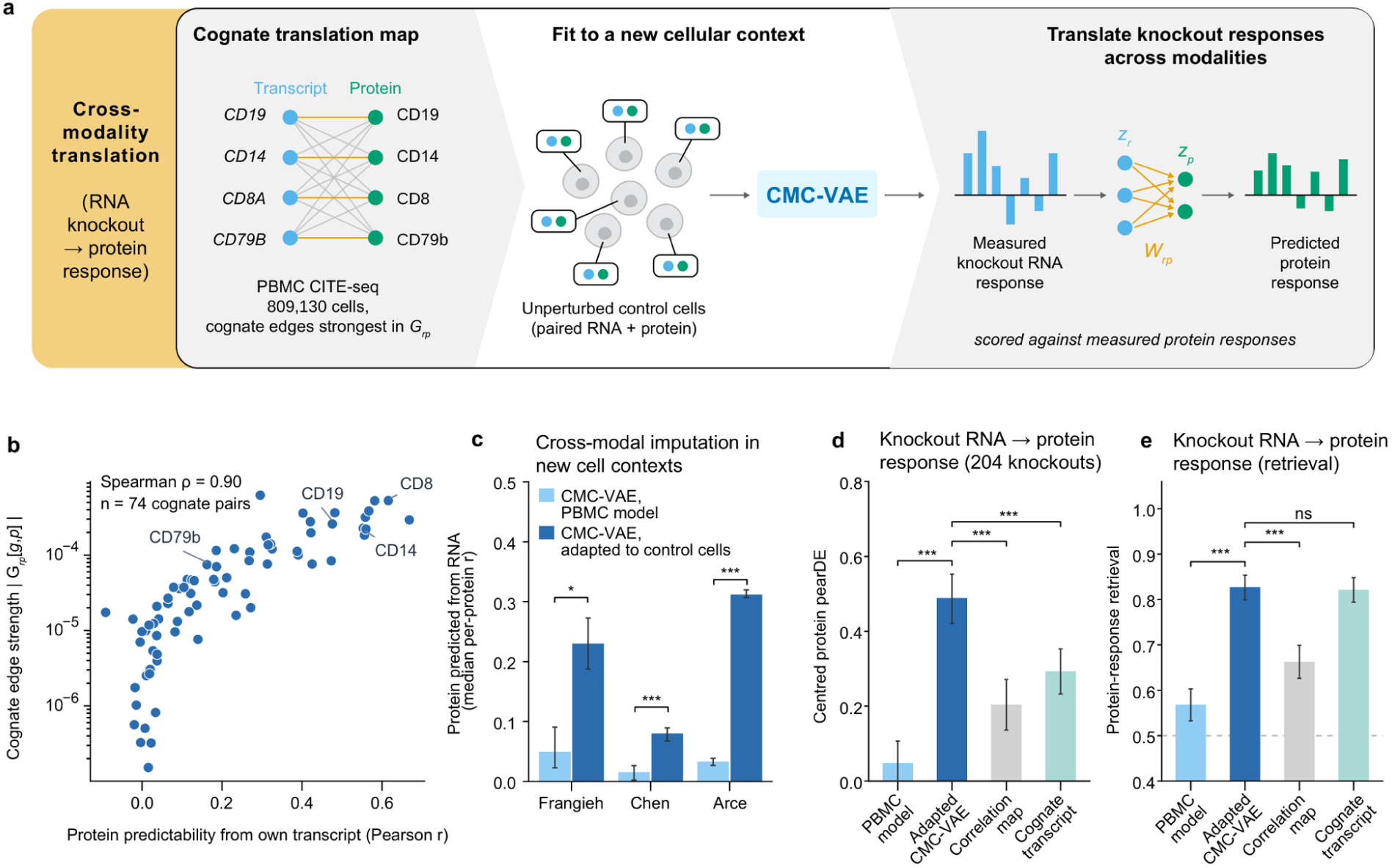
A cognate translation map and cross-modal translation of knockout responses. **a,** Schematic. CMC-VAE learns from paired PBMC CITE-seq (809,130 cells) an RNA-to-protein map whose strongest edges link transcripts to their cognate proteins (left). The model is then fitted to a new cellular context using only that context’s unperturbed control cells, with paired RNA and protein and no perturbation labels (middle). Each knockout’s measured RNA response is encoded, passed through the RNA-to-protein latent block W_rp_ and decoded into a predicted protein response, which is scored against the measured protein response (right). **b,** Cognate edge strength |G_rp_[g, p]| (log scale) against how well each protein is predicted by its own transcript in the training cells (Pearson’s r), for 74 cognate transcript–protein pairs. **c,** Protein imputation from RNA in control cells of each screen (median per-protein Pearson’s r) for the PBMC-trained model and after adapting its encoders, decoders and latent graph to the screen’s unperturbed control cells without perturbation labels (Methods). Bars show the mean of three models; error bars, range across models; brackets, two-sided Wilcoxon signed-rank test across proteins (seed-mean per-protein r; 7, 76 and 84 proteins in Frangieh, Chen and Arce) with Benjamini–Hochberg correction across screens. Ridge regression fitted on the same control cells reached 0.50, 0.15 and 0.36 (Results). **d,e,** Translation of each knockout’s measured RNA response into a predicted protein response, compared with the measured protein response (204 knockouts with at least three DE proteins: 53 Frangieh, where only 7 panel proteins are measured, 130 Chen and 21 Arce; metrics averaged over three models): mean-centred protein pearDE (d) and retrieval of the correct knockout (e; chance 0.5, dashed line). Comparators are an RNA–protein correlation map from the training corpus and the response of the knockout’s cognate transcripts (centred pearDE defined for 199 knockouts); ridge regression fitted on the same control cells reached centred pearDE 0.70 and retrieval 0.91 (Results). Bars show means with 95% bootstrap confidence intervals over knockouts; brackets compare the adapted model with each comparator (bootstrap over knockouts). *P < 0.05, ***P < 0.001; ns, not significant.

We next asked whether the model can convert RNA into protein in the CRISPR screens, which differ from the PBMC training corpus in cell type, culture and assay. Applied directly, the PBMC model imputed surface proteins from RNA poorly in screen control cells (median per-protein r = 0.05, 0.02 and 0.03 in Frangieh, Chen and Arce; Fig. 4c). We therefore adapted the encoders, decoders and latent graph to each screen’s unperturbed control cells, without any perturbation labels (Methods), which raised imputation to 0.23, 0.08 and 0.31 (Fig. 4c). We then used the adapted model to translate each knockout’s measured RNA response into a predicted protein response. Across 204 knockouts with at least three DE proteins (53 in Frangieh, 130 in Chen and 21 in Arce), adaptation increased agreement between predicted and measured protein responses from 0.05 to 0.49 (mean-centred pearDE; 95% confidence interval (CI) 0.42–0.55) and retrieval of the correct knockout from 0.57 to 0.83 (95% CI 0.80–0.85), with significant gains in each screen. The adapted model exceeded an RNA–protein correlation map from the training corpus (0.20 and 0.66; P < 10⁻⁴) and, in centred pearDE, the response of the knockout’s cognate transcripts alone (0.29; P = 2 × 10⁻⁴), but not in retrieval (0.83 versus 0.82; P = 0.75) (Fig. 4d,e). A ridge regression fitted on the same control cells was more accurate (imputation 0.50, 0.15 and 0.36; centred pearDE 0.70; retrieval 0.91), as expected for a supervised model fitted directly in each screen, however the adapted CMC-VAE recovers about 70% of the ridge centred pearDE while keeping an explicit cross-modal graph. The same architecture fitted from scratch on these control cells translated at least as well (Supplementary Note 15), so this result reflects the ability of the CMC-VAE architecture to learn a cross-modal map from a context’s paired control cells, rather than transfer of the PBMC map itself.

### Interventional fine-tuning learns perturbation responses

Finally, we asked whether interventions can refine the latent graph. We first tested this in simulation, where the true graph is known, in a best case that matches the model class: a planted linear latent structural model observed through a linear decoder, with perfect interventions on latent variables. Here, observational training left edge orientation close to chance, whereas fine-tuning the adjacency on one known-target intervention per latent variable raised orientation accuracy from 0.59 to 0.84 with 8 latent variables and from 0.49 to 0.75 with 16 (q ≤ 6 × 10⁻⁴; 15 replicates each) and improved prediction of held-out responses (pearDE from 0.52 to 0.83 and from 0.42 to 0.75; orientation in Fig. 5b). We then froze the encoder and decoder of the observational model and fine-tuned only the latent adjacency on the CRISPR knockouts pooled across the three CITE-seq screens, holding out entire genes for evaluation (Fig. 5a; Methods). Fine-tuning raised held-out RNA pearDE from 0.01 to 0.20 overall, but label-shuffled and screen-mean targets gave the same gains (0.20 for both; centred pearDE 0.055 with real versus 0.054 with label-shuffled targets), so on these screens the gain reflects the response shared across knockouts, and raw pearDE stayed below that of a model-free predictor using the mean response of the training knockouts (0.44; Supplementary Note 15 and Supplementary Fig. 10). Fine-tuning on these screens, including with acyclicity or asymmetry penalties, also left the RNA graph nearly symmetric (Supplementary Note 15).

**Fig. 5.**
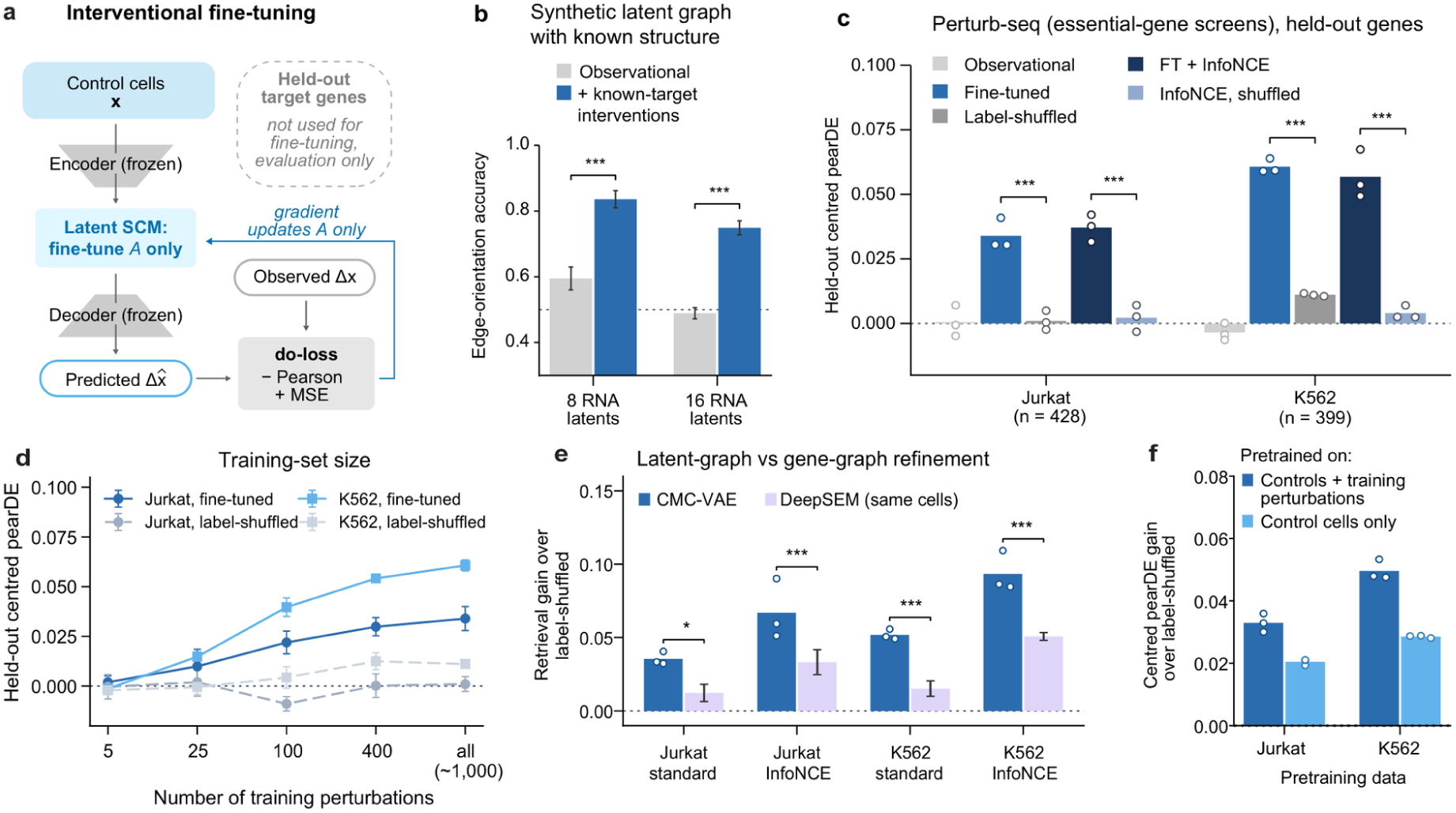
Interventions refine the latent graph and, with large-scale data, teach it gene-specific effects. **a,** Encoders and decoder remain fixed while the latent adjacency (gate logits and weights) is updated with a supervised response objective on training perturbations; evaluation uses held-out genes. **b,** Simulation with a planted latent graph observed through a linear decoder: edge-orientation accuracy of CMC-VAE after observational training and after fine-tuning the adjacency on one known-target intervention per latent variable (8 or 16 RNA latent variables; 15 replicates; mean ± s.e.m.; paired two-sided t-tests with Benjamini–Hochberg correction). **c,** Held-out mean-centred pearDE on the Jurkat and K562 essential-gene CRISPRi Perturb-seq screens (428 and 399 held-out genes) for the observational model and for fine-tuning with the standard and contrastive (InfoNCE) objectives, each with its label-shuffled control (mean of three draws); bars, mean of three models; dots, individual models; brackets, fine-tuned versus label-shuffled. **d,** Held-out mean-centred pearDE against the number of training perturbations (mean ± s.d.). **e,** Gain in held-out retrieval over the label-shuffled control for CMC-VAE (dots, individual models) and for DeepSEM trained and fine-tuned on the same cells with a mask of its 73,728 largest entries (the number of trainable latent-graph parameters of CMC-VAE), the same loss and stopping rule (mean ± s.d.); brackets, difference of gains. **f,** Gain in held-out centred pearDE over label-shuffled fine-tuning for models pretrained on control cells and training perturbations or on control cells only; dots, individual models. Tests in c and e use seed-averaged per-gene differences (bootstrap and two-sided Wilcoxon tests) with Benjamini–Hochberg correction within the panel. Dotted lines, zero or chance. *P < 0.05, **P < 0.01, ***P < 0.001.

Learning gene-specific effects required more perturbation data than the CITE-seq screens provide, which had at most 293 training units with modest responses dominated by the shared component. We therefore repeated the experiment on large-scale CRISPRi Perturb-seq screens of essential genes in K562^29^ and Jurkat^30^ cells, training an RNA-only CMC-VAE on control cells and the cells of training perturbations (without perturbation labels) and fine-tuning its latent graph on about 1,000 training perturbations, with 428 (Jurkat) and 399 (K562) held-out target genes and three independently trained models per cell line (Methods). Fine-tuning increased the perturbation-specific component of held-out responses (mean-centred pearDE from about 0 to 0.034 in Jurkat and 0.061 in K562), whereas label-shuffled fine-tuning did not (≤ 0.011; gain over shuffled +0.033 and +0.050, bootstrap P < 5 × 10⁻⁴, positive for every model; Fig. 5c). A contrastive objective (InfoNCE)^31^ increased held-out gene retrieval, the probability that a prediction matches its own knockout better than another held-out knockout, from 0.52 to 0.57 in Jurkat and from 0.51 to 0.61 in K562. The gene-specific gain appeared with 25–100 training perturbations and increased with the number of training perturbations (Fig. 5d). We compared this with DeepSEM, a gene-level structural model, trained and fine-tuned on exactly the same cells with an edge mask matching the 73,728 trainable latent-graph parameters of CMC-VAE, the same loss and the same stopping rule. CMC-VAE gained more retrieval over its label-shuffled control than DeepSEM in all four comparisons (both cell lines, standard and contrastive objectives; difference +0.02 to +0.04, q ≤ 0.02; Fig. 5e). Because every latent edge is shared by all genes through the decoder, whereas the out-edges of a held-out gene in DeepSEM are never updated, this comparison indicates that a shared latent parameterisation transfers to held-out genes more effectively than a gene-level edge set. The gain did not require perturbed cells during pretraining: models pretrained on control cells alone still improved over label-shuffled fine-tuning (centred pearDE +0.020 in Jurkat and +0.029 in K562, positive for every model; Fig. 5f).

These gene-specific effects are small and supervised predictors trained on the same perturbations were more accurate (Supplementary Fig. 11 and Supplementary Table 10). Interventional fine-tuning of the latent graph therefore learns gene-specific effects within an explicit structural model when perturbation data are sufficient.

## Discussion

CMC-VAE shows what paired single-cell multi-omics can and cannot reveal about causal regulation on its own. Without regulatory priors, trajectories or perturbation labels, the model learned a sparse, feature-resolved representation of RNA and surface-protein organization, including co-expression and immune-program structure and a transcript–protein map enriched for cognate pairs. CRISPR screen experiments showed that the model ranks knockout-responsive genes better than correlation and totalVI baselines in the development screens and that this ranking relies on the learned RNA graph; in a screen withheld from development it matched the correlation baseline, and no method predicted knockout-specific responses there. These experiments also exposed an asymmetry in what the model had learned. The observational RNA graph identified which genes were related but not which genes are regulators, whereas the cross-modal architecture, fitted to a new cellular context with unlabelled control cells, learned a map that translated knockout RNA responses into protein responses. Updating the graph with a small number of interventions improved held-out response correlation, mainly through the response shared across knockouts, and with large-scale perturbation data it learned small gene-specific effects for unseen genes, more so than a gene-level model fine-tuned on the same cells, though less accurately than supervised read-outs. Thus, observational data reveal the connections of the cross-modal network and a cognate transcript-to-protein map, whereas gene-specific perturbation effects require sufficient interventional data.

Most single-cell multi-omic methods are designed primarily to align modalities, impute missing measurements or compress shared variation into a common latent space, as exemplified by MOFA+, totalVI, MultiVI and more recent foundation-model approaches^7–9,32^. Other methods incorporate a supplied regulatory graph or temporal ordering to make cross-modal relationships more interpretable^10,33^. CMC-VAE places an explicit, editable graphical network between and within modalities and projects that structure directly onto measured genes and proteins. This change of objective entails a trade-off. totalVI was the stronger protein imputer, whereas CMC-VAE produced a substantially more concentrated cognate map and exposed the structural object used for intervention. Likewise, the recovered antigen-presentation, interferon and inflammatory programs are best viewed as biological validation and an interpretability aid, rather than as the discovery of novel pathways. Paired measurements on the same sample make these cross-modal comparisons possible without computationally matching different cells, but pairing by itself does not establish the direction of regulation.

An important limitation of observational only training is that even though the observational RNA graph was sparse, reproducible and useful for ranking perturbation-responsive genes, it did not resolve which gene regulated which. Moreover, recovery of regulon databases that are labelled as directed (DoRothEA, CollecTRI) was almost matched by an undirected correlation baseline, showing that agreement with a directed reference can be driven largely by co-expression. Two properties of the model explain this result. First, the reconstruction objective identifies the composite decoder–SCM map, allowing changes in the adjacency to be absorbed by the decoder. Second, within the linear-Gaussian regime that we assumed, members of a Markov equivalence class cannot be distinguished from their observational distribution^2,3^. Consistent with this, none of the variants we tested (for example, altered regularization, nonlinear structural equations or an acyclicity constraint) produced an orientation that was consistent across cohorts, and sparsity penalties on the latent graph made the network asymmetric without orienting it (Supplementary Table 11). Other assumptions, such as non-Gaussianity, environment shifts or temporal information, can help identify direction when they are satisfied,^5,34^ but we could not establish that these conditions hold in our setting, and the variants we tested did not orient the graph (Supplementary Notes 6 and 7). Therefore, the RNA graph should be interpreted as an undirected map of candidate regulatory interactions, not as an observationally identified causal regulatory network, and orientation should be treated as part of the problem that interventional data must supply, as our simulation illustrates (Fig. 5b).

The RNA-to-protein map was more specific. CMC-VAE assigned approximately eightfold larger weights to cognate than to non-cognate transcript–protein pairs, and cognate-edge strength closely tracked the extent to which each surface protein was predictable from its transcript. The architecture was also useful beyond the training corpus: fitted to a screen’s control cells without perturbation labels, it translated measured knockout RNA responses into protein responses better than a corpus correlation map and, in profile agreement, better than the cognate transcripts alone. A ridge regression fitted on the same control cells remained more accurate, so the value of the adapted model lies in combining useful translation with an explicit, inspectable cross-modal graph. The orientation of this map does not follow from a larger translation block relative to the feedback block, because that magnitude ratio changes with latent dimension and regularization, and the feedback block carries little independent receptor-to-RNA information after removing the transpose of the translation map. The translational map therefore captures cognate translation, but not post-transcriptional regulation or receptor signalling broadly.

Freezing the encoders and decoder preserves the observational representation, while updating the comparatively small structural component forces the interventional signal to correct edges instead of relearning cell state. On the CITE-seq screens, the gain in held-out RNA response correlation mainly reflected the response shared across knockouts in each screen, a property also described for other perturbation benchmarks.^35^ However, learning gene-specific effects required more perturbations. On large-scale Perturb-seq, the gene-specific gain emerged with about 100 training perturbations and grew thereafter, and exceeded that of DeepSEM fine-tuned on the same cells while not requiring perturbed cells during pretraining. This procedure is related to differentiable causal discovery from interventions^36^, but differs from perturbation-response models trained end to end to predict unseen treatments^37,38^. Recent benchmarks show that generalization to unseen perturbations and cellular contexts remains difficult and strongly context dependent, and complex models have been found to be no more accurate than simple baselines in several common benchmarks^39,40^, although models that incorporate curated biological priors have recently reported gains (for example, PIE).^13^ Here, our narrower objective is to adapt a network already learned from abundant paired observations. Consistent with these benchmarks, supervised predictors using the same gene representations outperformed fine-tuning of the latent graph on held-out genes. However, the responses we learned did not carry over between cohorts as a graph fine-tuned on two cohorts did not predict well on a held-out third without cohort-specific perturbation data, indicating that the effective edges depend on cellular composition, assay and perturbation context. Choosing which perturbations to measure could make this step more efficient.

This study has several limitations. First, training and validation are restricted to PBMC training data and immune-related perturbation settings, with 2,692 genes and a 144-marker surface-protein panel rather than a genome-wide proteome. Second, the low level of protein feature complexity potentially led to the worse model performance on the protein space than RNA space. The protein–protein block recovered some known surface complexes but did not outperform co-expression on the broader panel, which mainly reflects cell-type identity. Protein-level validation experiments were also uneven across cohorts, because the overlap with our antibody panel and the size of protein-level response differed widely, with the smallest held-out set containing only a handful of Arce perturbations. Cross-modal translation in a new cellular context also required fitting the model to that context’s control cells. Model selection used the three development screens, whereas baselines were run with default settings. Furthermore, our *in silico* interventions act on latent dimensions, not directly on genes, so they only approximate gene-level perturbations. The predicted response of this operation is dominated by its edge-scaling term, so predicted responses are more similar across knockouts than measured responses, and because it uses absolute decoder loadings it ranks response magnitude without fixing the sign of the target’s own change (Supplementary Fig. 12), however a signed knockdown shift corrects this and was used for Perturb-seq fine-tuning. The model also did not capture the effect of perturbation durations, dosage or cell-to-cell variation in perturbation efficiency. Finally, for the ranking task evaluated here, adding more observational data did not necessarily improve generalization as pooling more heterogeneous cohorts was less useful than selecting context-relevant data, even with the same number of cells. Hence, more data, even with diversity, does not always provide more information in causal learning, and selecting training data that matches the intended application may matter more than scale.

These findings suggest a general design principle for causal multi-omics. Abundant paired observations can be used to learn a stable representation, candidate interaction structure and modality-specific mappings, whereas targeted perturbations can be reserved for the smaller but indispensable task of learning perturbation-specific effects. Broader protein panels, including intracellular proteomics, paired RNA–chromatin, RNA–methylation or spatial measurements could extend the block structure of our causal framework, although the identifiability of the causal content must be reconsidered for each assay across the datasets rather than assumed from the architecture. Future models could allow the graph to capture context-dependent or hierarchical relationships in order to distinguish shared from cohort-specific regulatory mechanisms. On the other hand, time-resolved and combinatorial perturbations could test dynamics that a steady-state model cannot resolve. More broadly, our results suggest a practical way to build multi-modal causal models of the cell. Observational data show which genes and proteins are connected and recover a transcript–protein map enriched for cognate pairs. Perturbations can then supply what observation alone cannot, which is the effects of intervening on specific genes. An intervenable causal graph of this kind offers a possible correction layer for multimodal virtual-cell models, in which observational pretraining supplies coverage and perturbation experiments add interventional constraints, although supervised read-outs of the same representation currently predict perturbation responses more accurately than graph-based refinement.

## Methods

### The CMC-VAE model

For each cell, the RNA input contains 2,692 features and the protein input contains 144 features. Separate residual encoders produce diagonal Gaussian posteriors over exogenous latent variables: 128 dimensions for RNA and 64 for protein. We concatenate these variables into a 192-dimensional vector, ε, and regularize its posterior toward a standard normal prior. Encoder architectures and training settings are given in Supplementary Note 1.

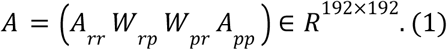

The latent adjacency, A, contains four blocks: RNA-to-RNA connections, RNA-to-protein connections, protein-to-RNA connections and protein-to-protein connections. We use the source-by-target convention, so entry *A_ji_* represents the coefficient from latent variable *j* to latent variable *i*. Each entry combines a learned gate with a real-valued weight, using the continuous gate relaxations described in Supplementary Note 1.^41–43^ Diagonal masks remove self-loops within the RNA and protein blocks. The endogenous latent variables satisfy

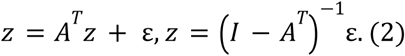

The spectral radius, ρ(A), is the largest absolute eigenvalue of A. When ρ(A) < 1, the inverse above exists and can be expressed as a convergent sum of powers of Aᵀ. We penalized spectral radii above 0.9 and measured a radius of approximately 0.89 in the reported final model. The penalty encourages stability but does not guarantee it at every optimization step. Nonlinear variants are listed in Supplementary Note 5 and Supplementary Table 4. The nonlinear variant used 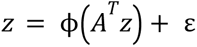 and fixed-point iteration; stability of that nonlinear system depends on the full update map.

#### Linear decoder and projected scores

Each modality has a linear decoder with a bias term. The RNA decoder maps the RNA latent variables to reconstructed RNA features, and the protein decoder maps the protein latent variables to reconstructed protein features. Let D be the block-diagonal matrix formed from the two decoder matrices. We summarize the latent adjacency in measured feature space using

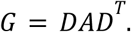

The four blocks of G provide scores for gene–gene, gene–protein, protein–gene and protein–protein pairs. For example, the RNA-to-protein score matrix is

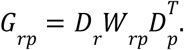

These scores combine latent connections with decoder loadings and are used for reference-network comparisons. They are not assumed to be independently identified direct effects between measured features.

#### Training objective

We minimized a regularized variational objective combining reconstruction error, a free-bits KL term and penalties on the model parameters. Reconstruction used mean squared error on the standardized RNA and protein inputs. The KL term regularized the diagonal Gaussian posteriors toward the standard normal prior; the free-bits thresholds limited the penalty applied to low-divergence latent dimensions.^44^ The structural terms penalized spectral radii above 0.9 and departures of the decoder columns from orthogonality. The final decoder also used a group-lasso penalty with 16 latent groups, as specified in Supplementary Note 1.

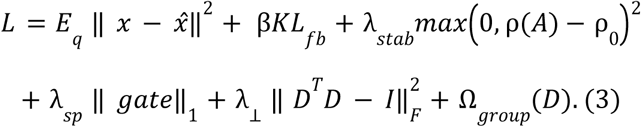

Here, ρ_0_ = 0. 9, *KL_fb_* denotes the free-bits divergence from the standard normal prior, and Ω*_group_* denotes the weighted decoder group-lasso term. Coefficients determine which penalties are active in each configuration.

The final model did not include an acyclicity penalty. We compared it with a variant that applied the DAGMA penalty to the RNA and protein diagonal blocks.^45^ This comparison constrains within-modality cycles; it does not by itself prohibit cycles passing through both modalities. The final configuration and the 27-configuration comparison are described in Supplementary Notes 1 and 5.

### Interpretation of the latent and projected networks

The projection G = DADᵀ is a deterministic summary of the fitted parameters. If D has orthonormal columns, the latent adjacency can be recovered from this summary and the known decoder:

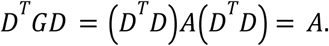

This identity concerns recovery of the latent matrix. It does not imply that individual feature-space entries preserve the support, sign or magnitude of individual latent edges. For a full-column-rank decoder, the corresponding inverse uses the pseudoinverse of D. Biological interpretation of the projected entries therefore requires external evaluation.

In an unrestricted linear decoder model, reconstruction alone can leave the decoder and structural model ambiguous. Writing B = D(I − Aᵀ)⁻¹, any alternative adjacency with an invertible structural operator can be paired with D′ = B(I − A′ᵀ) to preserve B. This construction need not preserve CMC-VAE’s block-diagonal decoder or its regularization constraints. It illustrates a potential factorization ambiguity rather than proving arbitrary reorientation within the implemented model class.

Classical causal-identifiability results impose additional assumptions. Linear Gaussian directed acyclic graphs with unrestricted disturbance variances can be observationally indistinguishable within a Markov equivalence class.^2,3^ Equal disturbance variances can identify a Gaussian DAG under the relevant assumptions, and LiNGAM identifies linear acyclic models with independent non-Gaussian disturbances and no unobserved confounding among the modeled variables.^4–6^ These results do not directly establish identifiability or non-identifiability of CMC-VAE’s learned cyclic latent representation. We therefore report the symmetry, perturbation predictions and control experiments of the fitted models as empirical findings.

An ideal intervention replaces the structural equation of the intervened variable. Changes can propagate to descendants, but an observed response need not identify a direct child, and cancellation or weak effects can obscure downstream changes. Our decoder-guided operation modifies distributed latent connections associated with a measured gene. Its usefulness is assessed by prediction of experimental responses; we do not assume that it is exactly equivalent to a gene-level do-intervention.

### Decoder-guided perturbation prediction

To estimate the response to perturbing gene *g*, we used its absolute RNA-decoder loadings to weight the RNA latent dimensions: *s* = _|_*D_r_* [*g*, *j*]_|_/*max_j_* _|_*D_r_* [*g*, *j*′]_|_, for *j* = 1, …, *l_r_*. We scaled the associated latent connections, solved the modified linear system with the same encoded exogenous input, and decoded the change:

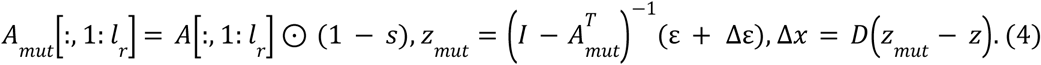

Here ε is set to the encoder posterior means of up to 4,000 control cells (a fixed random subsample), and the knocked-out gene is additionally clamped toward zero expression through an exogenous shift, Δε. This shift drives gene g’s exogenous input toward its zero (mean) value and propagates through the encoder weights; on the standardized scale, zero expression is the feature mean rather than the absence of signal. In practice, this shift contributed about 0.01% of the norm of the predicted response in the development screens, which is therefore dominated by the edge-scaling term (Supplementary Note 15). Δx̂ is the decoded expression difference averaged over those control cells with the knocked-out gene excluded from scoring. The vector (1 − *s*) is broadcast over the selected RNA columns. Under the source-by-target convention in equation (2), this expression scales contributions into the RNA latent variables. Because each gene is represented across several latent dimensions, this operation modifies a distributed representation rather than directly setting the measured gene’s expression to zero. It is an approximation whose usefulness is assessed against experimental responses, rather than an exact gene-level do-intervention. Because the edge scaling uses absolute loadings, this operation ranks the magnitude of responses but does not by itself fix their sign: the predicted change of the knocked-out gene was negative for 93%, 60% and 61% of knockouts in Frangieh, Chen and Arce (Supplementary Fig. 12). We therefore use it for zero-shot ranking and, for Perturb-seq fine-tuning, use a signed knockdown shift (Methods, ‘Large-scale Perturb-seq’).

Graph controls shuffled or removed connections within a selected block while retaining the fitted encoders and decoder. The RNA reversal control transposed the square block *A_rr_*.

### Data and preprocessing

The observational corpus comprised 809,130 paired CITE-seq profiles from the Hao healthy PBMC reference (161,764 cells from 8 donors) and the Stephenson COVID-19 atlas (647,366 cells from 130 donors).^15,16^ We retained the published cell calls and did not apply additional cell-level filtering. Median library sizes were 6,863 and 3,067, median numbers of detected genes were 2,026 and 1,241, and median mitochondrial fractions were 5.7% and 2.7%, respectively. Genes detected in fewer than three cells were removed before vocabulary construction.

RNA inputs were taken from the Hao SCT “counts” assay and the Stephenson “raw” layer. The training-cohort comparison also included COMBAT and Su (Supplementary Note 2).^46,47^ Corpus selection used structural and perturbation-prediction metrics. The CRISPR screens were not part of observational gradient training, but their performance informed this model-selection comparison. The Papalexi screen used for the final test was not used in any model development.

We constructed a vocabulary of 2,692 genes. We first selected 2,000 highly variable genes using the seurat_v3 criterion with cohort as the batch key, then removed mitochondrial and cytoplasmic ribosomal genes. We added CollecTRI transcription factors that were highly variable or detected in at least 5% of training cells, highly variable RNA-binding proteins annotated with GO:0003723, and perturbation-target genes detected in at least 5% of training cells. Thus, target identities and biological gene lists informed feature selection, although the final observational loss did not use regulatory edge labels or perturbation outcomes.

The protein vocabulary was the intersection of the two antibody panels after marker-name harmonization and removal of isotype controls, yielding 144 markers. RNA values were library-size normalized to 10,000 and log-transformed. Protein values were centred-log-ratio normalized within each cohort. Both modalities were standardized to zero mean and unit variance per feature at the encoder input. These per-feature standardization statistics (mean and variance) were computed across all cells before the train/validation/test split; recomputing them on the training cells alone changed per-feature means by a median of 0.0004 s.d. and standard deviations by 0.08%, and perturbation-screen cells never entered these statistics.

We used a fixed 80/10/10 split of the pooled cells for training, validation and testing (seed 42). This was a cell-level split and therefore does not by itself test generalization to unseen donors. We used cohort-aware highly variable gene selection without explicit ComBat correction. Cohort silhouette scores ranged from 0.08 to 0.19 in the examined data and latent representations, and donor groups were mixed under the reported diagnostic. Per-gene excess kurtosis remained approximately unchanged after ComBat correction (Supplementary Note 13). These diagnostics assess the particular batch correction and embedding summaries tested; they do not exclude all technical contributions to the observed variation.

#### Perturbation cohorts

We evaluated the Frangieh Perturb-CITE-seq screen (174 evaluated perturbation units; 7 overlapping proteins), the Chen FOXP3-regulator Perturb-icCITE-seq screen (122 units; 78 proteins) and the Arce CRISPRi Perturb-CITE-seq cohort (23 units; 84 proteins).^22–24^ Singlet cells were processed using RNA library-size normalization and log transformation and protein centred-log-ratio normalization. Knockout cells carried a single targeting guide and control cells only non-targeting guides; in the Frangieh data, 18,292 of the 34,611 cells previously labelled as controls also carried a targeting guide and were excluded. The Papalexi ECCITE-seq screen in IFN-γ-stimulated THP-1 cells^25^ (18 targets in the model vocabulary with at least 30 cells; 2,386 non-targeting control cells) was processed in the same way. Predictions were scored on RNA and protein features shared with the model vocabulary. These counts describe the final evaluation population and differ from the threshold-screening denominators in Supplementary Note 11.

### Evaluation

#### Differential expression

DE was defined by condition-matched pseudobulk analysis. For each screen we formed pseudobulks of at least 10 cells per guide and stratum (mean log-normalized RNA and CLR protein), with strata defined by culture arm (Frangieh: control, IFN-γ and co-culture), donor (Chen) or hashtag-defined activation and cell state (Arce). Each knockout was compared with non-targeting pseudobulks from the same strata using a linear model y ∼ knockout + stratum, empirical-Bayes variance moderation (limma-trend-like) and two-sided moderated t-tests, with Benjamini–Hochberg correction within each knockout. A feature was called DE when q < 0.05 and |LFC| ≥ 0.10 (RNA) or 0.05 (protein), where LFC is the stratum-adjusted difference in mean log-normalized expression (or CLR) between perturbed and control cells. Perturbations with fewer than 30 cells were excluded. Model-development analyses (training-cohort and configuration comparisons; Supplementary Notes 2, 5–8, 11 and 14) used a cell-level definition (two-sided Mann–Whitney U P < 0.05 without multiple-testing correction, with the same LFC floors, against all control cells pooled per screen, including for Frangieh the uncorrected control set), which is also used for the single-condition Papalexi and Perturb-seq screens; the development conclusions were unchanged with Benjamini–Hochberg-corrected DE (Supplementary Note 11), and the Papalexi and Perturb-seq conclusions with Benjamini–Hochberg-corrected or pseudobulk DE (Supplementary Note 16). For in silico prediction, the same set of up to 4,000 control cells (a fixed random subsample) was encoded for every knockout. Signed pearDE was computed only when at least five features were DE, and AUROC and the protein read-out required at least three DE features.

#### Prediction metrics

AUROC measured how well absolute predicted changes ranked DE features above non-DE features. Signed pearDE was the Pearson correlation, across DE features, between the predicted change (in the standardized decoder-output space) and the measured log-fold change (on the stored log-normalized or CLR layer). It measures agreement between response profiles, rather than sign accuracy or the orientation of individual causal edges. Metrics were calculated for each eligible perturbation unit and summarized within the cohort. Measured effects are differences in mean log-normalized expression (or CLR) between perturbed and control cells; for brevity we refer to them as log-fold changes (LFC).

#### Shared-response references

Because knockouts in the same screen share a common response component, we report three additional quantities (Supplementary Note 15). The mean-response reference predicts, for each knockout, the mean measured response of the other (or, for held-out evaluation, the training) knockouts in the same screen. Mean-centred pearDE subtracts this mean response from both the predicted and the measured response before computing pearDE. Held-out gene retrieval is the probability that a knockout’s prediction correlates more strongly with its own measured response than with that of another held-out knockout, computed on features that were DE in at least one training knockout and excluding all target genes (chance 0.5).

#### Untouched screen

The Papalexi screen was evaluated once, with the three final models of each method and a frozen pipeline, after its end points, comparators and tests had been written down. The pre-specified primary end points were mean-centred pearDE and retrieval, which did not differ from zero or chance (Supplementary Note 15); RNA-response AUROC was computed exactly as for the development screens. Knockouts with at least three DE genes (13 of 18) were evaluable, and comparisons used paired two-sided t-tests across knockouts with Benjamini–Hochberg correction.

#### Reference networks

We scored absolute projected interaction weights against binarized co-expression, STRING, DoRothEA and CollecTRI references.^17–19^ OmniPath was used in the equal-latent-dimension comparison.^48^ Reference-network AUROC measures agreement with known relationships. Forward-to-reverse weight asymmetry and correlation with the transposed score matrix were used as descriptive summaries of network symmetry.

#### Cognate null models

Cognate enrichment was the median absolute RNA-to-protein score for the 74 transcript–protein pairs divided by the median non-cognate score. We recomputed enrichment under 1,000 randomizations for each of three nulls: permuting entries of the projected RNA-to-protein score matrix G_rp_, permuting protein-column assignments, and a column-norm-preserving null. We report the distance of the observed enrichment from the null mean in units of the null standard deviation. None of the 1,000 randomized values under any null reached the observed enrichment. Preserving column norms does not by itself preserve graph degree.

### Interventional fine-tuning

For fine-tuning, we split target genes between training and evaluation sets, keeping each gene in the same split across all three cohorts. Of 418 perturbation units representing 401 genes, 280 genes were assigned to training (293 units: 127 Frangieh, 150 Chen and 16 Arce) and 121 disjoint genes to evaluation (125 units: 54 Frangieh, 64 Chen and 7 Arce). Fine-tuning updated the entire latent SCM, all four gate-logit and all four edge-weight blocks, with the encoders and decoder frozen. For each held-in knockout (excluding the knocked-out gene), the target was the observed log-fold change over differentially expressed features and the prediction was the decoded soft-knockout response over the same features; the direction term used the raw LFC and the magnitude term used a standard-deviation-scaled LFC. The protein term L_rp_ was the analogous negative-Pearson-plus-squared-error loss on protein DE features (weight 0.5), and a spectral-radius guard kept ρ(A) < 0.92. Each step used the encoder posterior means of 3,000 control cells per cohort, averaged. The three seeds (42, 1 and 2) are three independently trained observational models, each with its own training-cell split, fine-tuned with the same gene split and hyperparameters. With the condition-matched ground truth, 87 held-out units had a defined RNA pearDE (51 Frangieh, 31 Chen and 5 Arce) and 38 a defined protein pearDE.

The supervised response objective combined a negative Pearson-correlation term, a squared-error term, a protein-response term and a spectral-stability penalty:

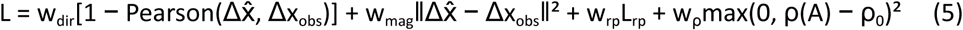

In equation (5), the observed response is the raw log-fold change in the direction (Pearson) term and the per-feature standard-deviation-scaled log-fold change in the magnitude term; predictions are in the standardized decoder-output space. The direction, magnitude, protein-response and stability terms had weights 1, 1, 0.5 and 0.1, respectively and the spectral-radius target was 0.9. Fine-tuning used a learning rate of 5 × 10⁻⁴, 150 steps and 3,000 cells. Frozen weights were checked before and after training. Evaluation used the disjoint target genes. Supplementary Note 14 reports the leave-one-cohort-out experiment. The fine-tuning weights were chosen during development with held-out-gene performance in view; because the label-shuffled and screen-mean controls use identical weights, this choice does not affect the control comparisons.

#### Control fine-tuning

Control runs used the same protocol with the targets replaced by (i) the response of a different, randomly assigned training knockout (label shuffle, three draws), (ii) the mean response of the training knockouts in the cohort (mean target), (iii) sign-flipped targets or (iv) targets centred on that mean response. A contrastive variant added an InfoNCE term^31^ (temperature 0.1) that rewards a higher correlation of each prediction with its own target than with the targets of the other knockouts in the batch; its weight was chosen on an inner 30% split of the training genes.

#### Large-scale Perturb-seq

We used the essential-gene CRISPRi Perturb-seq screens in K562^29^ and Jurkat^30^ cells (log-normalized expression; 3,000 features shared by both lines, comprising 1,685 perturbation targets and 1,315 highly variable genes). Target genes were split 70/30 (seed 42), identically in both lines, and DE features were defined against each line’s non-targeting cells by the cell-level criterion above. An RNA-only CMC-VAE with the same latent structural model (192 latent variables, so that its 73,728 trainable latent-graph parameters are a gate and a weight for each of the 192² latent positions) was trained on non-targeting control cells and the cells of training perturbations, or on control cells only, with three models (seeds 42, 1 and 2) per line and setting. Fine-tuning used the protocol above with a signed knockdown shift (the minimum-norm change of the exogenous input that sets the decoded expression of the target to zero), the same control targets and a number of steps chosen on an inner 30% split of training genes. DeepSEM was trained on the same cells as each CMC-VAE model and fine-tuned with a mask of its 73,728 largest-magnitude entries (the number of trainable latent-graph parameters of CMC-VAE; about 0.8% of the gene–gene matrix, one parameter per edge) under the same loss, objectives, label-shuffled arms and stopping rule. Ridge and k-nearest-neighbour baselines mapped each target’s CMC-VAE decoder loadings or its co-expression profile in control cells to its response, fitted on training perturbations, and reduced-rank ridge regression constrained predictions to the span of the decoder. GEARS,^37^ the linear model of Ahlmann-Eltze et al.^40^ and ridge regression on scGPT gene embeddings^32^ were scored with the same scorer on the same held-out genes. Comparisons used seed-averaged per-gene differences with bootstrap confidence intervals and two-sided Wilcoxon tests, with Benjamini–Hochberg correction within the panel. Details are given in Supplementary Note 16.

Control-cell adaptation. For each CITE-seq screen and model, the encoders, decoders and latent graph were adapted on 90% of the screen’s unperturbed control cells, using the observational objective with an added cross-modal reconstruction term, and early stopping on the remaining 10%. No perturbation labels were used, and the procedure was specified before evaluation. Imputation was scored as the median per-protein Pearson correlation between imputed and measured protein in control cells.

Translation was scored by encoding each knockout’s measured RNA response and decoding a protein response, which was compared with the measured protein response by mean-centred pearDE and retrieval (204 knockouts across the three screens; metrics averaged over three models). Comparators were an RNA–protein correlation map from the training corpus, the response of the knockout’s cognate transcripts and ridge regression from RNA to protein fitted on the same control cells. Confidence intervals and P values come from bootstrap resampling of knockouts.

### Baselines and synthetic study

We trained totalVI on the same observational cells (batch key, donor; 50 epochs). Because totalVI has no structural adjacency, its RNA-to-RNA and RNA-to-protein networks were the Pearson correlations between totalVI-denoised RNA and, respectively, denoised RNA (symmetric) and denoised protein, computed over a 60,000-cell training subsample. For an in silico knockout of gene g, the RNA and protein read-outs were the negated rows for g in these two correlation matrices (knockdown sign); protein imputation was evaluated on held-out cells with protein masked.^7^ GRNBoost2 used the CollecTRI transcription-factor list.^20^ DeepSEM was trained with an L1 penalty (α = 100) on its gene-by-gene structural adjacency W and spectrally stabilized by renormalizing W after each epoch to a spectral radius ρ(W) ≤ 0.9; responses propagated through DeepSEM’s own (I − Wᵀ)⁻¹.^21^ DeepSEM knockout responses were read out with the knockdown sign, as for CMC-VAE. For the matched DeepSEM fine-tuning, the network was frozen except for a trainable-edge mask on the 73,728 largest-magnitude entries of W (equal to the number of trainable latent-graph parameters of CMC-VAE), updated with the same response loss, knockdown operator, step budget, early-stopping rule, spectral guard, scoring and held-out gene split as CMC-VAE. Matched refers to the shared loss, do-operation and held-out genes; the trainable parameters are the top-magnitude edges of DeepSEM rather than the full latent SCM of CMC-VAE. The models used a common evaluation procedure on the same cohort collection. Comparisons were aligned by perturbation unit where specified below.

For the synthetic comparison, we generated data from a planted sparse acyclic graph with controlled cross-modal connections. Exogenous variables followed Gaussian, Laplace or Student’s t distributions with eight degrees of freedom. They were propagated through the structural model and a random linear decoder. We compared the projected read-out with DirectLiNGAM applied to the latent variables and to variables obtained by pseudo-inverting the decoder.^6^ These settings distinguish access to the latent variables from evaluation of a learned measurement model. Results are reported in Supplementary Note 6 and Supplementary Table 3. For interventional recovery (Fig. 5b), CMC-VAE was fitted to planted block structural models (8 or 16 RNA and 4 or 8 protein latent variables) observed through a marker-structured linear decoder, and only A was then fine-tuned with the response loss of equation (5) on one perfect intervention per RNA latent variable (do(z_k_) = −2 s.d.; 2,000 cells per knockout), evaluated on the other marker gene of each latent variable; 15 replicates per latent size.

### Statistics and reproducibility

Perturbation metrics in Figure 3 are reported as mean ± s.e.m. across evaluated perturbation units. In Figure 3b, brackets report the more conservative of a joint bootstrap over knockouts and control cells (999 replicates, DE recomputed in each replicate) and a paired two-sided t-test on aligned per-perturbation values (unpaired for the correlation baseline), with Benjamini–Hochberg correction across all Figure 3 comparisons; in Figure 3d, joint-bootstrap P values were adjusted within the panel. Figure 3c used paired two-sided t-tests across knockouts with Benjamini–Hochberg correction. In Figure 4c, imputation by the adapted and PBMC models was compared per protein with two-sided Wilcoxon signed-rank tests with Benjamini–Hochberg correction across screens. Figure 4d,e report means with 95% bootstrap confidence intervals over knockouts, and comparisons use bootstrap P values. Figure 5b used paired two-sided t-tests across simulation replicates; Figure 5c,e used seed-averaged per-gene differences with bootstrap and two-sided Wilcoxon tests. P values in Figure 5 were Benjamini–Hochberg-adjusted within each panel.

Figure 5c, e and f show the mean of three independently trained models (seeds 42, 1 and 2) per cell line and setting, with individual CMC-VAE models as dots; Figure 5d shows mean ± s.d. across models and random training-gene subsets. These seed-level error bars summarize a different source of variation from the per-perturbation tests.

Cognate score distributions were compared using a one-sided Mann–Whitney U test and the randomized nulls described above. Surface-complex recovery was assessed by permutation of edge labels. Residual receptor-target AUROCs in Supplementary Figure 7b were compared with chance by target-label permutation tests at P < 0.05. Final-model seed variation is reported in Supplementary Table 8 and Supplementary Note 12.

## Data availability

The Hao PBMC reference is available from Gene Expression Omnibus under accession GSE164378, the Stephenson atlas from ArrayExpress under E-MTAB-10026 and the Frangieh screen from the Single Cell Portal under SCP1064.^15,16,22^ The COMBAT and Su datasets are available from the repositories cited in their original publications.^46,47^ The Chen Perturb-icCITE-seq data are available from the DDBJ Sequence Read Archive under PRJDB16517, with processed data at Zenodo (doi:10.5281/zenodo.16020737).23 The Arce Perturb-CITE-seq data are available under GEO SuperSeries GSE271090, subseries GSE278572.^24^ Reference networks were obtained from CollecTRI, DoRothEA, STRING and OmniPath. The Papalexi ECCITE-seq screen^25^ is available from GEO under GSE153056. The K562^29^ and Jurkat^30^ Perturb-seq screens are available from Figshare+ (doi:10.25452/figshare.plus.20029387) and GEO (GSE264667).

## Code availability

The CMC-VAE code, trained model weights and scripts to reproduce all analyses and figures are available at https://github.com/goodarzilab/CrossModalCausal-VAE.

## Acknowledgements

HG is a Core Investigator, and this research was funded by Arc Institute. We thank Chiara Ricci-Tam (Arc Institute) for help with editing the figures.

*Use of AI tools.* Large language models including Claude Code and ChatGPT were used to assist with language editing, structuring of manuscript sections and formatting of references and figure legends. All AI-assisted text was reviewed and revised by the authors, who verified all scientific content, analyses, statistics and citations and took full responsibility for the manuscript. Furthermore, Claude Code was used to generate the code and analyses for this manuscript, however the creative and critical direction of the paper was driven by the authors on this paper.

## Author contributions

S.G. conceived the study, developed the method, performed the analyses and wrote the manuscript. V.N. contributed analyses and helped with manuscript editing. N.L. and H.G. supervised the study and edited the manuscript. All authors approved the final manuscript.

