## Supplementary Information for "Multi-omics causal deep learning reveals regulatory structure while interventions refine perturbation predictions"

**Contents:** Supplementary Notes 1–16, Supplementary Figs. 1–12, Supplementary Tables 1–11 and Supplementary References. Development analyses (Supplementary Notes 2, 5–8, 11 and 14, the perturbation comparisons in Supplementary Note 1, Supplementary Figs. 4b, 5 and 6, Supplementary Tables 1, 4–6 and 8 and the RNA AUROC column of Supplementary Table 11) used the cell-level DE definition and, for Frangieh, the uncorrected control set (Methods); they guided model development and do not enter the main-figure comparisons.

#### Supplementary Note 1: Model and training details

Encoders are residual multilayer perceptrons (RNA, 2,692–512–256–128; protein, 144–128–64–64) with batch normalization, LeakyReLU activations and 10% input-feature dropout. The decoder is linear, LD-VAE style with a bias term. In the final model, gates use a Gumbel-sigmoid relaxation (temperature annealed from 1.0 to 0.5 at epoch 30; deterministic sigmoid at evaluation); hard-concrete $L_0$ gates[1] were available as a variant but not used for the reported model. The final model uses $\beta = 2.0$, free bits of 1.0 (RNA) and 0.5 (protein), no acyclicity constraint (cycles permitted), a spectral-stability weight of 20.0 ($\rho_0 = 0.9$), decoder group-lasso sparsity (16 latent groups, $\lambda_{RNA} = 1,000$), an orthogonality weight of 5.0, no knowledge prior loss, and latent dimensions of 128 (RNA) and 64 (protein) with mean-squared-error reconstruction loss. It was trained for 120 epochs with Adam on a single H100 GPU. The group lasso partitioned the 128 RNA and 64 protein decoder columns each into 16 contiguous groups (eight RNA or four protein columns per group); the penalty was the per-feature sparse-group norm, the sum over features and groups of the L2 norm of each feature’s decoder weights within a group, divided by the number of groups, with λ = 1,000 for RNA and 18 for protein, each scaled by a 0.05 ramp (effective weights 50 and 0.9); no gate L1 penalty was applied in the final model. The group-lasso decoder produced a projected GRN with a high fraction of near-zero weights (99% of edges below 0.1% of the maximum) and improved both perturbation prediction and regulon recovery relative to a feature-norm decoder.

Compared with an otherwise identical model using the DAGMA acyclicity penalty, allowing cycles had little effect on the learned observational structure, while improving TF-regulon recovery (DoRothEA AUROC from 0.64 to 0.68) and improved held-out perturbation prediction in the development evaluation (mean RNA-to-RNA AUROC from 0.65 to 0.67; cell-level DE). This is a modeling comparison on these metrics and does not establish that the fitted cycles correspond to biological feedback. Three training seeds (42, 1 and 2) reproduce the key metrics (Supplementary Table 8).

#### Supplementary Note 2: Training-cohort sweep

We compared training on Hao, Stephenson, COMBAT and Su, individually and in combinations.[2,3,4,5] Supplementary Table 1 lists all 15 non-empty cohort combinations and two additional final-pipeline entries.

Adding cohorts did not consistently improve RNA-response ranking. In the reported all-cell runs, Su alone achieved a mean AUROC of 0.669, while the four-cohort pool achieved 0.629. These comparisons show that a larger pooled dataset did not necessarily improve this prediction task. They do not isolate a single cause of the difference because cohort composition, feature vocabularies and other processing choices also require consideration.

The final Hao–Stephenson model used a different vocabulary pipeline from the corresponding sweep entry and scored 0.667 rather than 0.635 on mean RNA-response AUROC. That cohort was selected based on performance across multiple structural and perturbation metrics, including a reported mean protein pearDE of approximately 0.34 (development evaluation with the uncorrected Frangieh controls) and CollecTRI AUROC of 0.63. The same CRISPR screens were used during this comparison; their role in model selection and final evaluation is specified in Methods.

#### Supplementary Note 3: Immune program annotations

Each RNA latent was annotated by preranked gene set enrichment analysis (GSEA; gseapy) of its decoder loadings over the 2,692 genes against MSigDB Hallmark gene sets (minimum set size 5, maximum 500, 200 permutations), keeping the top pathway per latent at a false discovery rate (FDR) of $q < 0.1$; this annotated 31 of the 128 RNA latents of the final model. The strongest $A_{rr}$ edges among these labelled latents formed a program graph dominated by an antigen-presentation hub with interferon, TCR and JAK–STAT satellites (Supplementary Fig. 1a,b). In the projected RNA network, edge reciprocity was approximately 0.99, compared with approximately 0.01 under the degree-preserving null.[6] Four of the 50 genes with the largest outgoing weight sums were transcription factors. Outgoing weight sum correlated with decoder-row norm (Pearson's $r = 0.57$; Supplementary Fig. 1c,d), indicating that gene-level score magnitude partly reflects the decoder loadings.

When the same architecture was trained separately on healthy and on COVID-19 cells, the learned immune programs differed between the healthy and COVID-19 training groups (Supplementary Fig. 2). The healthy model was dominated by a single antigen-presentation hub (20 of 36 significant latents), whereas the disease model halved that hub (10 latents) and expanded the type I and type II interferon programs (IFN-α/β, from 3 to 9 latents; IFN-γ, from 3 to 6) and apoptosis (from 2 to 5), consistent with the interferon-driven signature of severe COVID-19 (one fit per condition, without seed replication). Differences in cohort and cell-type composition, as well as latent annotation and fitting variability, limit their interpretation as disease-specific regulatory changes.

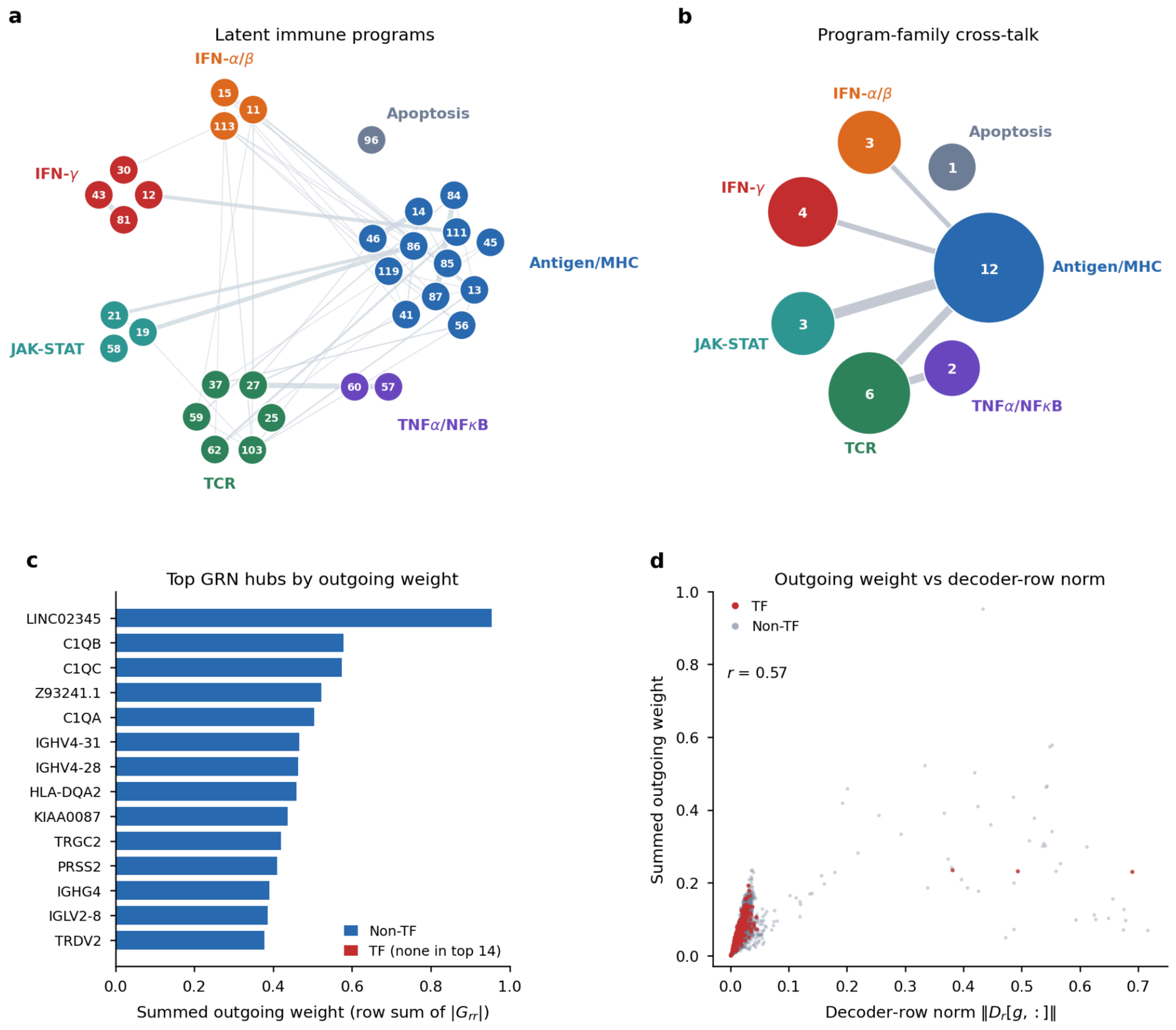


**Supplementary Fig. 1 | Immune-program annotations and RNA-network weight structure. a**, The 31 RNA latent variables with immune-program enrichment at FDR < 0.1, grouped by program family. Edges show the selected strongest latent connections and line width represents connection magnitude. Numbers identify latent dimensions. **b**, Summed absolute connection weights between program families; node area reflects the number of annotated latent dimensions. **c**, The 14 genes with the largest outgoing absolute weight sums in the projected RNA network; none of these 14 is a transcription factor. The statistic of four transcription factors among the top 50 genes in Supplementary Note 3 uses the larger set. d, Outgoing weight sum (the total absolute projected RNA-network weight leaving a gene, summed over its targets; a weighted strength, not an edge count) versus RNA decoder-row norm for the evaluated genes; r denotes Pearson correlation. Gene-set definitions, edge selection and enrichment testing are described in Supplementary Note 3.

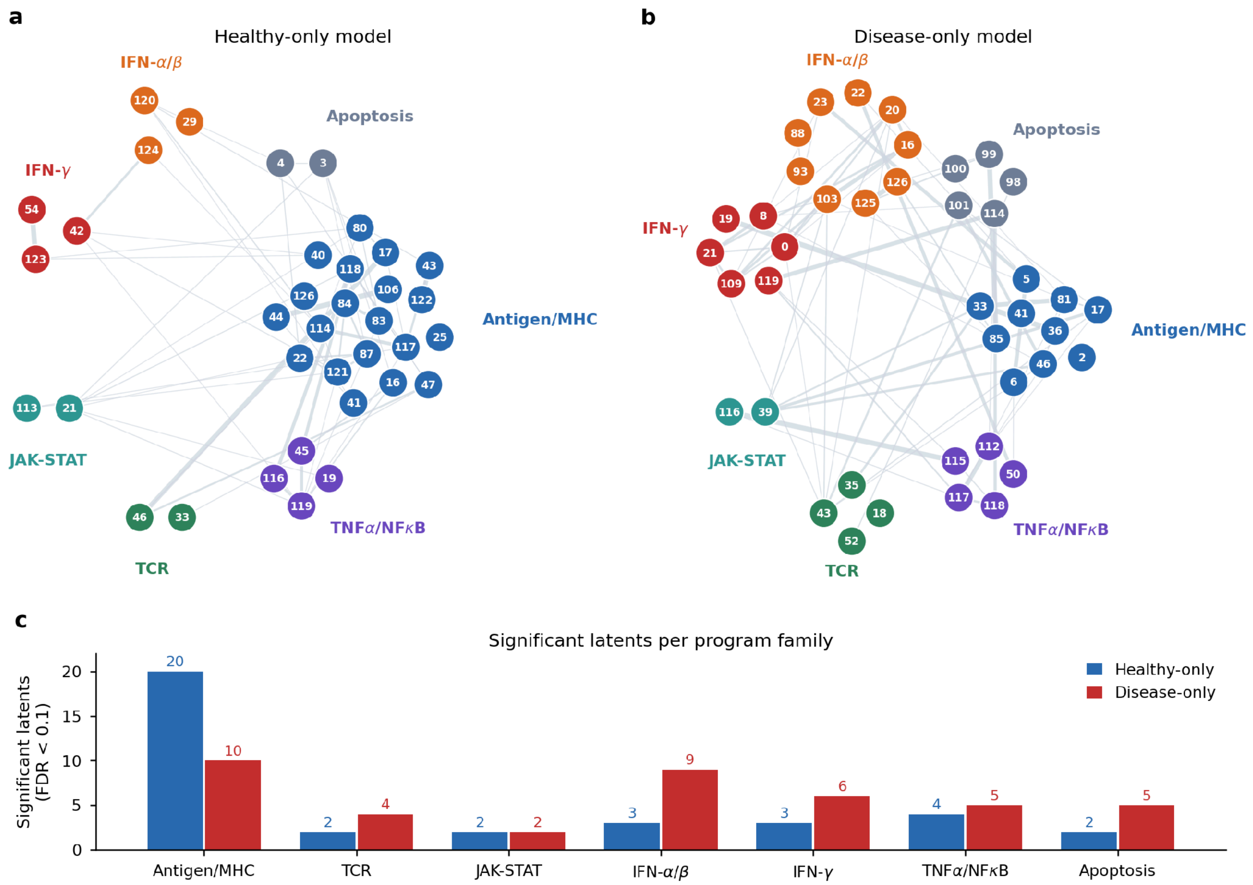


**Supplementary Fig. 2 | Program annotations in models trained on healthy and COVID-19 cells. a,b,** RNA latent-program graphs for the separately trained healthy model (36 annotated latents) and COVID-19 model (41 annotated latents). Colors indicate program families and edges show the selected latent connections. **c**, Number of latent variables assigned to each program family at FDR < 0.1. These counts describe independently fitted representations and are not aligned one-to-one latent variables. Cell composition, enrichment procedures and training settings are described in Supplementary Note 3.

#### Supplementary Note 4: External baselines and the protein network

On the reported protein-imputation evaluation, mean per-protein Pearson correlation was 0.502 for totalVI and 0.403 for CMC-VAE, compared with approximately 0.20 for the single-transcript baseline (Supplementary Fig. 3a and Supplementary Table 2).[7] CMC-VAE produced a more concentrated cognate map and higher scores on the DoRothEA and CollecTRI references than the learned-model baselines. However, the correlation baseline achieved 0.67 and 0.62 on those references, close to CMC-VAE's 0.68 and 0.63. These comparisons support recovery of known relationships but do not establish directed regulatory inference. The particular totalVI RNA read-out used here was symmetric by construction while GRNBoost2 used a supplied TF list. Neither observation establishes a general impossibility result for the methods. sciPENN was not included because the available implementation could not be run in the study's H100 environment.[8]

The protein network scored near chance against the examined intracellular interaction references. On the 36 curated surface-complex edges represented in the panel, its AUROC was 0.62, compared with

0.61 for marginal co-expression. For 15 direct physical connections, the scores were 0.71 and 0.68. The model did not consistently improve on co-expression across the full set or after the examined lineage adjustments (Supplementary Fig. 3c). The protein block was also less sparse by the chosen relative-weight threshold than the RNA and cross-modal blocks (Supplementary Fig. 3d,e).

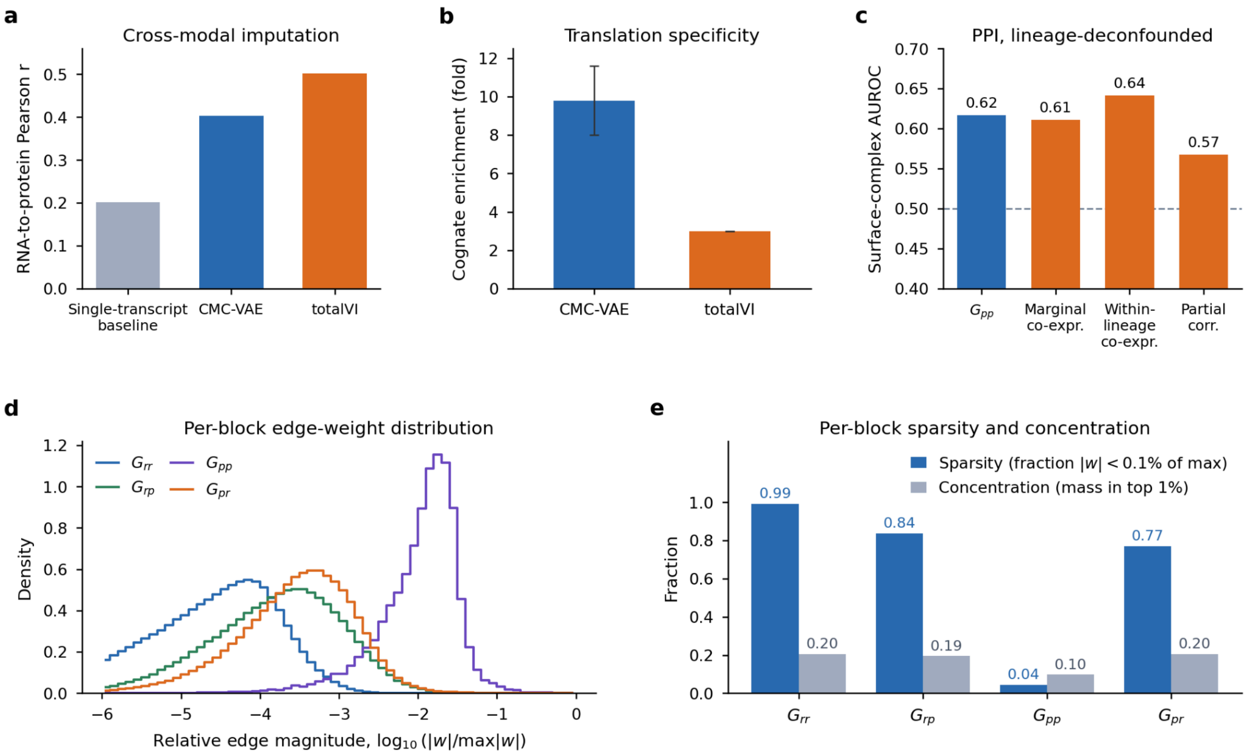


**Supplementary Fig. 3 | Protein imputation and the distribution of projected interaction scores. a**, Per-protein imputation correlations for the single-transcript baseline, CMC-VAE and totalVI. **b**, Cognate enrichment across three training seeds, shown as mean ± s.d. (9.8 ± 1.8); the deployed final-checkpoint value on the 74-pair set is 8.2 (main text). **c**, Recovery of 36 curated surface-complex connections using the protein map and the specified co-expression or partial-correlation controls. **d**, Distribution of edge magnitudes relative to each block's maximum absolute weight. **e**, Fraction of weights below 0.1% of that maximum and fraction of total absolute weight in the largest 1% of edges. The single-transcript model is a baseline.

### Supplementary Note 5: Architecture and regularization comparison

Supplementary Table 4 reports 27 configurations evaluated in Frangieh. The sweep varied KL weight, free-bits thresholds, exogenous-noise bottlenecks, decoder constraints, nonlinear structural equations, latent dimensions, initialization and directed-prior weight. Its baseline used a feature-norm decoder and differs from the final group-lasso configuration. RNA pearDE ranged from −0.117 to 0.086 in the listed configurations, and none of these regularizer changes produced signed RNA-response prediction.

Stronger exogenous-noise bottlenecks reduced the reported correlation between exogenous and endogenous latents from approximately 0.98 to 0.28, while reducing co-expression recovery and cognate enrichment. Changes to KL pressure had less effect on that correlation. Strong directed priors increased forward-to-reverse asymmetry, reaching approximately 798 at the largest listed weight, but did not improve RNA pearDE. Co-expression AUROC fell from approximately 0.94 to 0.75 in the high-prior settings (Supplementary Fig. 4).

A signed prior from KnockTF was also evaluated using 19 held-out transcription factors with perturbation data.[9] RNA pearDE changed from −0.06 to −0.09, with reported sign agreement of 0.57 and forward-to-reverse asymmetry of 1.13. These experiments show that the tested priors and regularizers did not improve response prediction on the evaluated targets. Increased asymmetry alone was not evidence of improved causal inference.

#### Supplementary Note 6: Synthetic orientation comparison

In the synthetic orientation study, Gaussian exogenous noise was unorientable by all methods, whereas Laplace and Student's *t* noise were perfectly oriented by DirectLiNGAM, both on the latent and after decoder inversion, but not by our second-order read-out (Supplementary Fig. 4a and Supplementary Table 3). In the trained model, ε was strongly non-Gaussian (median excess kurtosis 10.9; 99% of dimensions reject normality), yet the DirectLiNGAM causal ordering of the learned $z$ agreed with $A_{rr}$ only at chance (order agreement 0.46; chance ≈ 0.5), and the inferred ordering was itself only moderately reproducible across bootstraps (Kendall's τ ≈ 0.46 between bootstrap orderings) and did not improve perturbation direction. Marginal non-Gaussianity also does not establish LiNGAM's independence and acyclicity assumptions.[10]

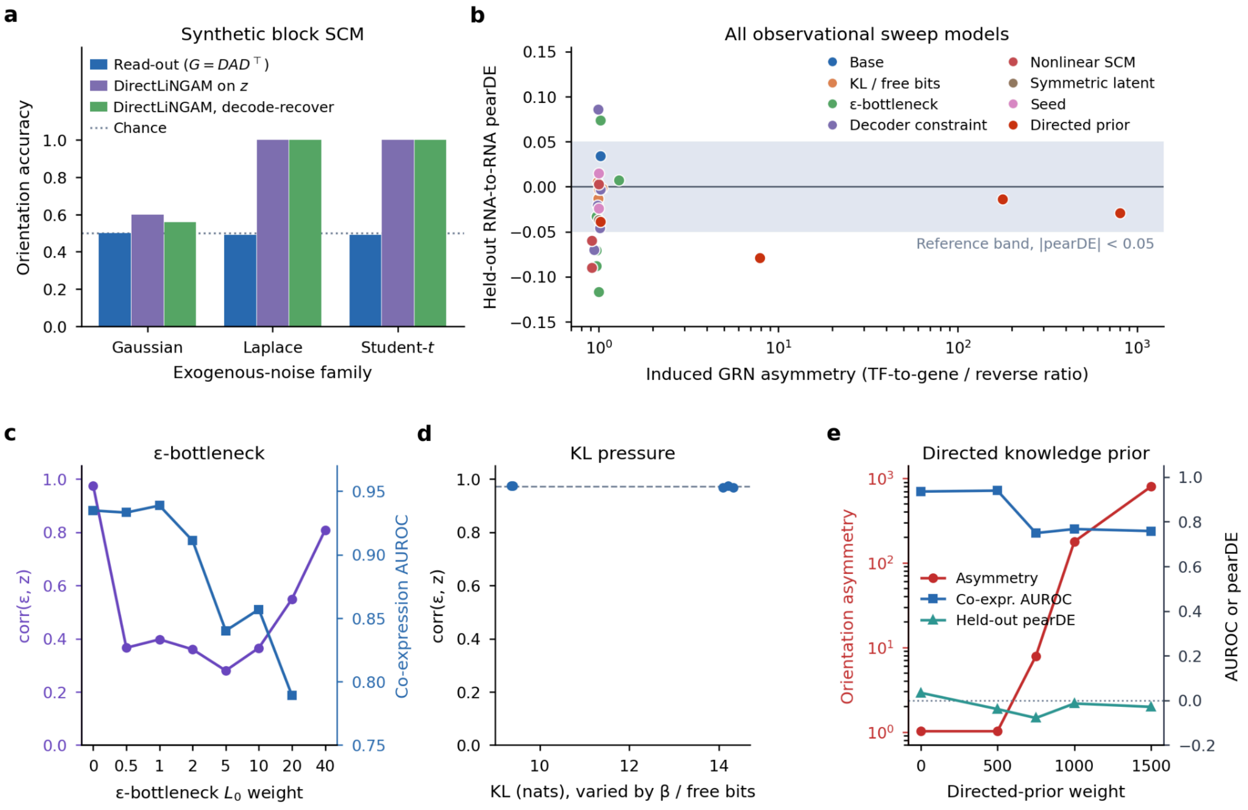


**Supplementary Fig. 4 | No observational lever recovers RNA-to-RNA direction.** **a**, Orientation accuracy on a synthetic block SCM for each exogenous-noise family, using our second-order read-out ($G = DAD^{\top}$), DirectLiNGAM on $z$, and DirectLiNGAM after decoder inversion; dotted line, chance. **b**, Held-out RNA-to-RNA pearDE of every model in the 27-configuration observational sweep against the induced GRN asymmetry (TF-to-gene/reverse ratio), coloured by lever; shaded band, |pearDE| < 0.05. **c**, $\varepsilon$-bottleneck: correlation between $\varepsilon$ and $z$ (left axis) and co-expression recovery AUROC (right axis) as a function of $L_0$ weight. **d**, KL pressure (varied by β and free bits) does not change corr($\varepsilon$, $z$). **e**, Directed knowledge prior: orientation asymmetry (left, log scale), co-expression AUROC and held-out RNA-to-RNA pearDE (right) as a function of prior weight.

#### Supplementary Note 7: Additional priors and training objectives

We examined additional training objectives motivated by representation-identifiability work: an auxiliary-variable-conditioned prior, a Laplace exogenous prior, total-correlation and HSIC dependence penalties, and an edge-reversal objective.[10–13] Auxiliary variables were cell type, donor and disease state. These variants test particular implementations of the ideas but they do not establish that the assumptions of the corresponding identification theorems hold for the data. We also tested whether an edge-reversal objective, supervised with measured perturbation outcomes, changes held-out RNA pearDE.

The Laplace, total-correlation, HSIC and reversal-objective variants produced projected RNA networks with forward–reverse correlations of at least 0.98 and small forward-minus-reverse pearDE differences (Supplementary Table 5 and Supplementary Fig. 5). Conditional priors reduced network symmetry, but their response-prediction changes differed across cohorts. The donor and disease priors also reduced Frangieh protein pearDE (development evaluation with the uncorrected controls) from 0.69 to −0.10 and 0.15, respectively (Supplementary Fig. 6a–d). Breaking symmetry therefore did not reliably improve the measured predictions.

We summarized the fitted linear operator using the normalized squared distance $\| M_{rr} - I\|_F^2 / \| M_{rr}\|_F^2$, where $M = \left(I - A^T\right)^{-1}$, of its RNA block from the identity and the maximum absolute RNA-to-protein weight. The first statistic was 77% in the final model and 9–34% in the conditional-prior variants; maximum cross-modal weight was also lower in those variants (Supplementary Fig. 6e,f and Supplementary Table 6). Conditioning only the RNA prior reduced some of the protein-prediction loss, but did not consistently improve RNA responses across cohorts. These quantities describe changes in the fitted operators; they are not direct estimates of biological variance explained.

To examine the shape of the learned exogenous distribution, we whitened the representation and compared marginal excess kurtosis along coordinate axes and random directions. The reported random-direction-to-coordinate ratio was 0.61 for the joint representation and 0.30 for RNA alone, compared with 1.02 for simulated isotropic Student's t noise and 0.024 for independent Laplace noise at dimension 128. This diagnostic suggests that heavy tails are not confined to the coordinate axes. It does not establish a precise elliptical fraction or demonstrate that all potentially identifiable information is absent.

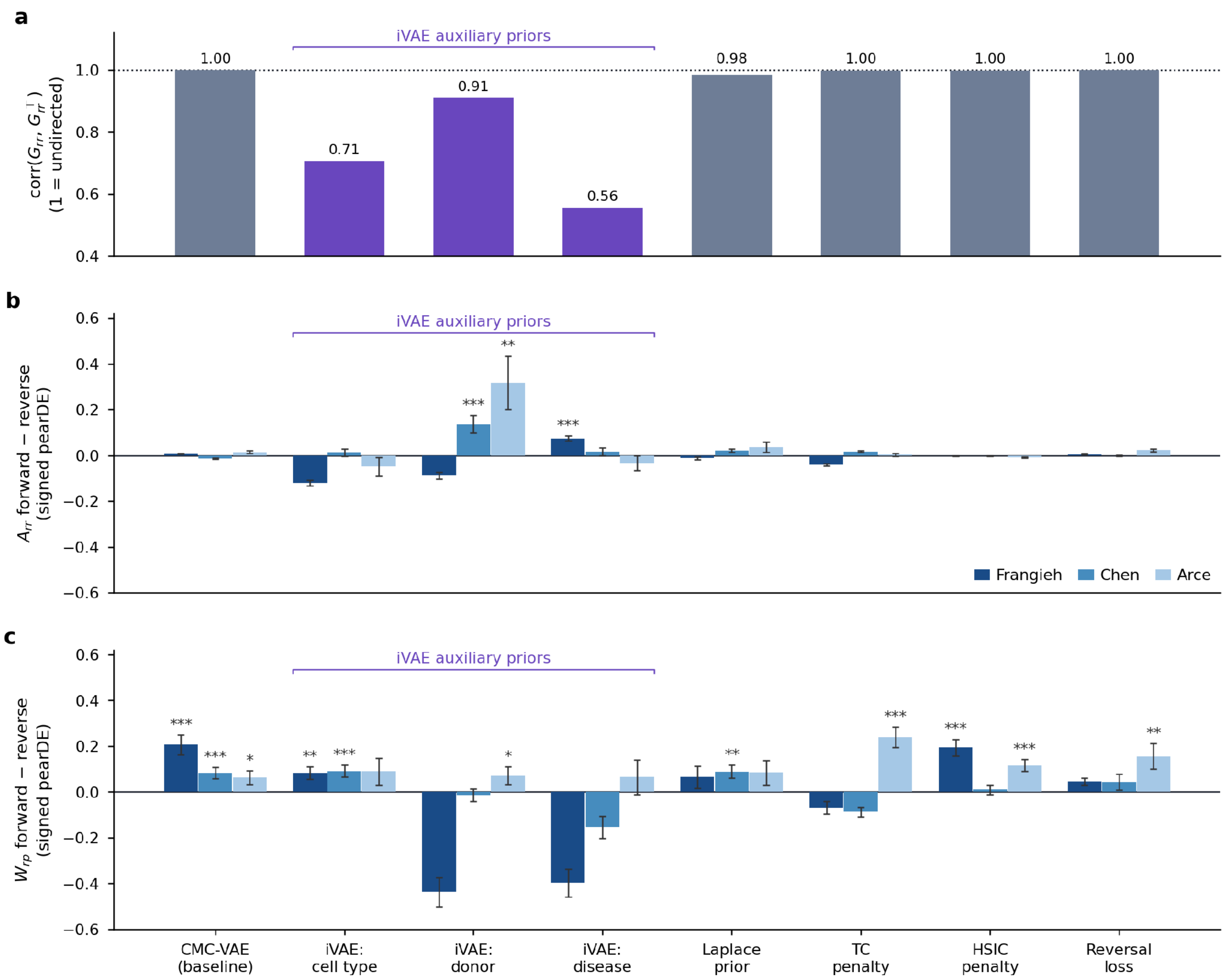


**Supplementary Fig. 5 | Graph symmetry and response prediction under additional training objectives. a**, Correlation between the projected RNA score matrix and its transpose; a value of 1 indicates symmetric scores. **b**, Difference in RNA pearDE between the learned and reversed square RNA block. **c, Difference in protein pearDE between the learned RNA-to-protein block Wrp and its replacement by the transposed protein-to-RNA block.** Values are shown separately for Frangieh, Chen and Arce. Error bars summarize s.e.m. across evaluated perturbation units with a defined pearDE (≥5 DE features): RNA panels n = 181, 191 and 21 and protein panels n = 55, 133 and 21 for Frangieh, Chen and Arce, respectively. Asterisks, paired one-sided *t*-test (forward > reverse), shown where |gap| ≥ 0.05: $^*P < 0.05$, $^{**}P < 0.01$, $^{***}P < 0.001$.

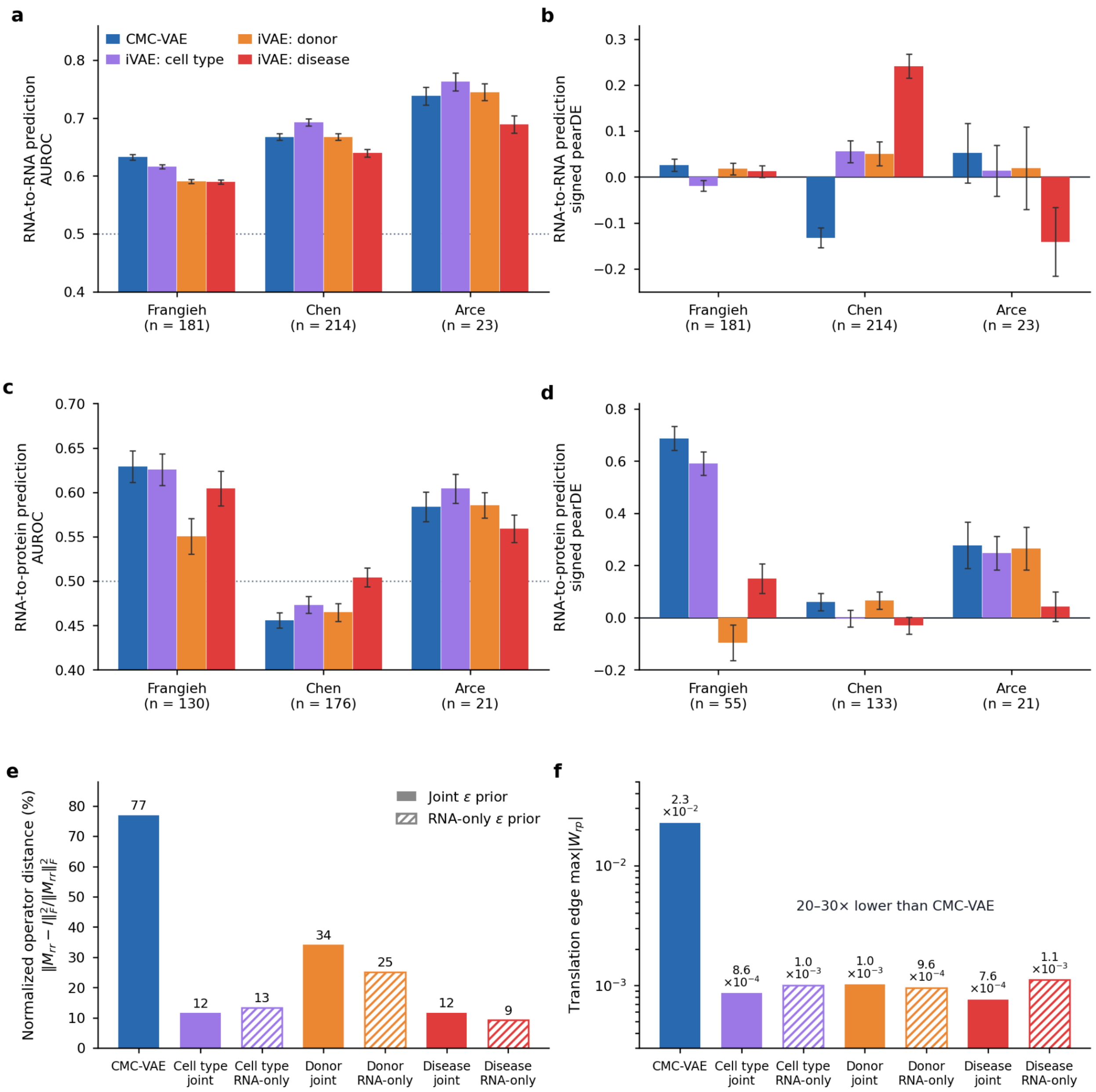


**Supplementary Fig. 6 | Prediction and fitted-operator changes under conditional priors. a–d, RNA and protein AUROC and pearDE for the final model and models with priors conditioned on cell type, donor or disease state (development evaluation: cell-level DE with the uncorrected Frangieh controls; the Frangieh protein pearDE of the final model is not supported by the corrected analysis, Supplementary Note 15).** Bars summarize eligible target units using mean ± s.e.m. **e**, Normalized squared distance of the RNA block of the resolvent from the identity, defined in Supplementary Note 7 and multiplied by 100 in the plotted percentage scale. **f**, Maximum absolute RNA-to-protein latent weight. Solid and hatched bars distinguish joint and RNA-only conditioning. The panel e value is 77% for the final model. Panels e,f are single-model parameter summaries, not fractions of biological variance explained.

### Supplementary Note 8: Equal latent dimensions

Giving the protein latent the same dimension as the RNA latent (128/128) left the GRN essentially unchanged (co-expression AUROC = 0.94; CollecTRI, 0.62; DoRothEA, 0.66) and balanced the cross-modal magnitudes (translation/feedback ratio 0.53), suggesting that the larger feedback weights in the 128/64 model are partly explained by the difference in latent dimensions. It did not improve recovery against intracellular PPI databases (OmniPath AUROC = 0.45), and the drop in RNA-response AUROC when the RNA latent block was shuffled was, if anything, larger (0.109). We therefore make no magnitude-based claim about cross-modal direction.

#### Supplementary Note 9: Per-protein translation decodability

We next examined how cognate translation strength varied across individual proteins. Across 74 cognate pairs, absolute RNA-to-protein edge strength correlated with the measured RNA–protein correlation (Spearman's ρ = 0.90). The totalVI-derived map showed a similar relationship (ρ = 0.86). Within-protein cognate specificity also tracked RNA–protein correlation in CMC-VAE (ρ = 0.82), and the cognate transcript ranked among the top approximately 10 of 2,692 inputs for some of the most strongly associated proteins.

Eighty-two percent of the cognate scores were positive. Negative scores included CD4, IL2RA, CXCR4 and CD62L-associated pairs, but their signs alone do not identify a post-transcriptional mechanism. Among non-cognate connections, reported precision at 10 for the evaluated lineage annotations was 0.33 for B cell proteins and 0.39 for monocyte proteins, compared with 0.004 under the specified null. The same analysis did not show comparable enrichment for T and NK cell proteins.

#### Supplementary Note 10: Inferred protein-to-RNA map

The protein-to-RNA map correlated with the transpose of the RNA-to-protein map (Pearson's r = 0.71). The fitted transpose component accounted for approximately half of the reported squared norm of the protein-to-RNA map (Supplementary Fig. 7a). We then evaluated the residual against the receptor-associated target sets shown in Supplementary Fig. 7b.

Residual scores rarely exceeded protein–gene co-expression for those sets. Immediate-early genes, including FOS, JUN, EGR1 and NR4A-family genes, received high scores across receptor rows and the prominent CD3-associated result was part of this broader pattern. These analyses provide little evidence for receptor-specific feedback in the fitted map.

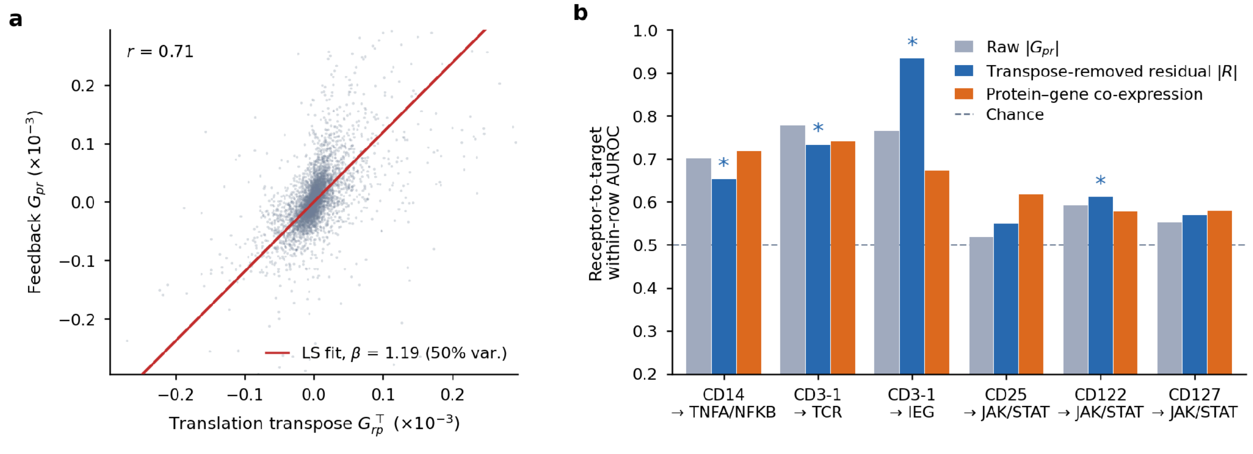


**Supplementary Fig. 7 | Shared structure in the two cross-modal maps. a**, Protein-to-RNA scores versus the transposed RNA-to-protein scores, with the fitted regression line and Pearson correlation. The displayed score scaling is indicated on the axes. **b**,

AUROC for six receptor→target-set rows (CD14→TNF/NF-κB, CD3→TCR, CD3→immediate-early genes, and CD25, CD122 and CD127→JAK–STAT), each excluding the receptor's own cognate transcript and ranked over the target genes present in the model vocabulary (approximately 31), using the original protein-to-RNA scores, the residual after removing the fitted transpose component, and protein–gene co-expression. A horizontal line marks AUROC = 0.5. The residual rarely exceeds co-expression (for example, CD3→TCR: raw 0.78, residual 0.73, co-expression 0.74). Asterisks indicate residual AUROCs above chance in a target-label permutation test (5,000 permutations, $P < 0.05$); above-chance does not test improvement over co-expression. The residual analysis is described in Supplementary Note 10.

#### Supplementary Note 11: Differential-expression outcome sensitivity

The primary differential-expression definition and effect-size scale are specified in Methods. Two counts appear in this work and are not interchangeable. The final evaluation counts of 174 (Frangieh), 122 (Chen) and 23 (Arce) perturbation units under the condition-matched definition (181, 214 and 23 under the cell-level definition used in this note), which are used throughout the main text, figures and Supplementary Table 7, are the perturbations whose target gene is present in the model feature vocabulary and that have at least three differentially expressed RNA genes. The larger screening denominators of 240 (Chen) and 25 (Arce) count all perturbations that pass the minimum-cell threshold with at least one differentially expressed gene, and are used only for the threshold-sensitivity analysis below.

In the reported threshold comparison, raising the RNA effect threshold from 0.10 to 0.25 left a median of 152 DE genes in Frangieh but a median of zero in Chen, where only 12 of 240 perturbations retained at least five DE genes. A protein threshold of 0.05 retained at least three DE proteins for 136 of 181 Frangieh, 182 of 240 Chen and 22 of 25 Arce perturbations. These are threshold-screening counts, not the evaluation counts (174, 122 and 23 under the condition-matched definition; 181, 214 and 23 under the cell-level definition). The Chen and Arce denominators of 240 and 25 therefore remain appropriate for this threshold analysis.

With DE features defined by Benjamini–Hochberg-adjusted $P < 0.05$ instead of the raw P value, 134 Frangieh, 106 Chen and 23 Arce perturbations remained eligible (42, 36 and 7 held-out units for fine-tuning). Conclusions were unchanged: observational RNA-response AUROC was 0.64 and protein pearDE 0.24, fine-tuning increased held-out RNA pearDE (for example, from −0.02 to 0.30 for the seed-1 model), and mean-centred pearDE remained close to zero, as with the primary definition.

#### Supplementary Note 12: Variation across seeds for training

We also checked whether the learned networks were consistent across random seeds 42, 1, and 2. The reported Spearman correlation of projected RNA scores on the defined shared-edge set was 0.75, and the top-1% edge Jaccard index was 0.38. Cognate RNA–protein scores had a Spearman correlation of 0.85. Furthermore, CLEC12A, CD36, CD19, CD14 and CD22 appeared among the strongest cognate pairs across runs. Non-cognate RNA-to-protein scores were less reproducible, with a reported correlation of 0.43. Supplementary Table 8 summarizes variation in the main metrics.

#### Supplementary Note 13: Data quality control

We compared cohort and donor mixing in the representations described in Methods and performed perturbation scoring within each cohort. Per-gene excess kurtosis of the input data had a median of approximately 17 and was the same for the two-cohort and ComBat-corrected inputs, showing that this

ComBat batch correction did not remove the heavy tails; it does not isolate whether the tails arise from biological heterogeneity, sparsity, normalization or other technical factors (Methods).

#### Supplementary Note 14: Interventional fine-tuning, additional analyses

The main reported fine-tuning experiment separated targets by gene while retaining all three cohorts in the training and evaluation sets. It therefore evaluates response prediction for other target genes within the represented cohort collection. It does not establish transfer to a cohort that supplied no perturbation data during training or model selection.

In the leave-one-cohort-out variant, the model was fine-tuned on two cohorts and evaluated on the third without cohort-specific interventions. RNA pearDE did not improve meaningfully in any held-out cohort: the change was −0.02 for Chen, −0.11 for Frangieh and −0.36 for Arce. These results show limited cross-cohort transfer under the tested procedure and mainly support the reported task of predicting new target genes within the cohorts represented during fine-tuning. Repeating this analysis with the control harness of Supplementary Note 15 for three independently trained models gave RNA pearDE changes of −0.05 (Frangieh), +0.02 (Chen) and −0.14 (Arce), with similar changes for label-shuffled targets; with a relaxed spectral guard ($\rho \leq 0.95$) the changes were −0.11, −0.05 and −0.02. The shared response was cohort-specific: the mean response of the other cohorts correlated with each held-out cohort's responses at −0.06 to −0.10, whereas the cohort's own mean response of the other knockouts correlated at 0.57–0.60. (These analyses used the cell-level DE definition of model development; Methods.)

#### Supplementary Note 15: Shared-response controls for perturbation metrics

Knockouts in the same screen share a common response component, for example responses to Cas9 cutting, guide delivery, selection or culture, so a high response correlation does not by itself show that a model has learned the effect of a particular gene.[14] We therefore compared every perturbation metric with four references computed from the same data: (i) a model-free mean-response predictor, the mean measured response of the other knockouts in the same screen (training knockouts only for held-out evaluation); (ii) mean-centred pearDE, the Pearson correlation between predicted and measured responses after subtracting that mean response from both; (iii) target-shuffled pearDE, in which each prediction is scored against a different knockout's measured response; and (iv) held-out gene retrieval, the probability that a knockout's prediction correlates more strongly with its own measured response than with that of another held-out knockout, computed on features that were DE in at least one training knockout and excluding all target genes (chance 0.5).

Observational predictions (Supplementary Figs. 8 and 9). With the condition-matched ground truth (Methods), signed RNA pearDE of all models remained close to zero. Protein-response AUROC of CMC-VAE was 0.49, 0.46 and 0.58 (Frangieh, Chen and Arce) and signed protein pearDE 0.05, 0.13 and 0.08; the measured protein panels changed little under the screened perturbations. Under the earlier cell-level evaluation, Frangieh protein pearDE had been 0.69, but that value reflected a control group that included cells carrying targeting guides and is not supported by the corrected analysis. Rankings by each gene's control-cell mean or s.d., which use no perturbation information, reached RNA-response AUROC of 0.72–0.93 (Supplementary Fig. 9), indicating that responsive genes share expression features across knockouts; the contribution of the learned network is therefore assessed with the graph ablations of Fig. 3d. Transposing $A_{rr}$ changed RNA-response AUROC by at most 0.019.

Untouched screen. On the Papalexi screen (13 knockouts, three models), the pre-specified primary end points did not differ from zero or chance: mean-centred pearDE 0.07 (95% CI −0.07 to 0.21; $P = 0.35$) and retrieval 0.53 (0.42–0.65; $P = 0.62$), similar to the correlation baseline, totalVI and DeepSEM ($q \geq 0.52$). RNA-response AUROC was 0.70 ($P = 6 \times 10^{-9}$ versus 0.5), above totalVI ($q = 2 \times 10^{-4}$) and similar to correlation and DeepSEM (Fig. 3c).

**Seed replication and adaptation control.** Repeating the Fig. 3 analyses for three independently trained models gave RNA-response AUROC of 0.610 ± 0.006, 0.664 ± 0.001 and 0.754 ± 0.007 (Frangieh, Chen and Arce; mean ± s.d.). Every significant Fig. 3b contrast kept its sign and $P < 0.05$ in each model, shuffling $A_{rr}$ lowered AUROC in every screen and model, zeroing $A_{rr}$ lowered it in Arce for two of three models (seed-mean $\Delta = 0.042$, $q = 0.014$), and CMC-VAE exceeded the correlation baseline for 83–84% of the 319 knockouts in every model. For cross-modal translation (Fig. 4d,e), the same architecture trained from random initialisation on the same control cells, with the same objective, data split, step budget and stopping rule, reached centred pearDE 0.62 and retrieval 0.89, compared with 0.51 and 0.84 for the adapted PBMC model on the same scorer (three models each), and similar control-cell imputation; the translation result therefore reflects the architecture fitted to each context rather than transfer of the PBMC map.

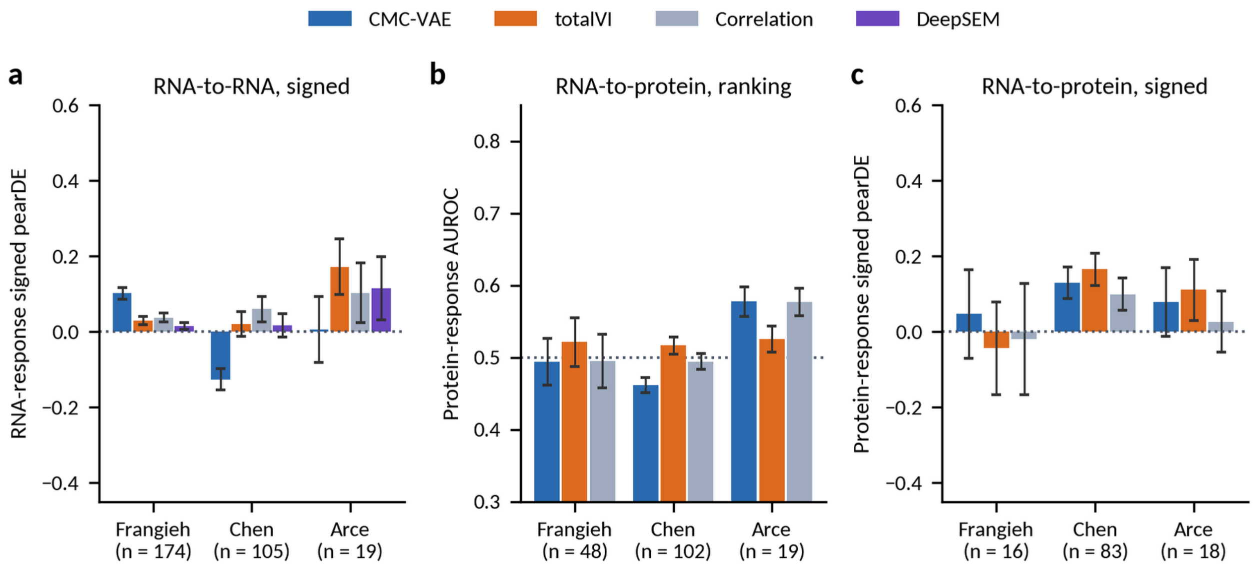


**Supplementary Fig. 8 | Signed and protein-response metrics for zero-shot prediction. a,** Signed RNA-response pearDE in the three development screens for CMC-VAE, totalVI, the correlation baseline and DeepSEM. **b,c,** Protein-response AUROC (b) and signed protein pearDE (c) for models with a protein read-out. Bars, mean ± s.e.m. across knockouts; ground truth as in Fig. 3; n under each screen. Dotted lines, AUROC = 0.5 or zero.

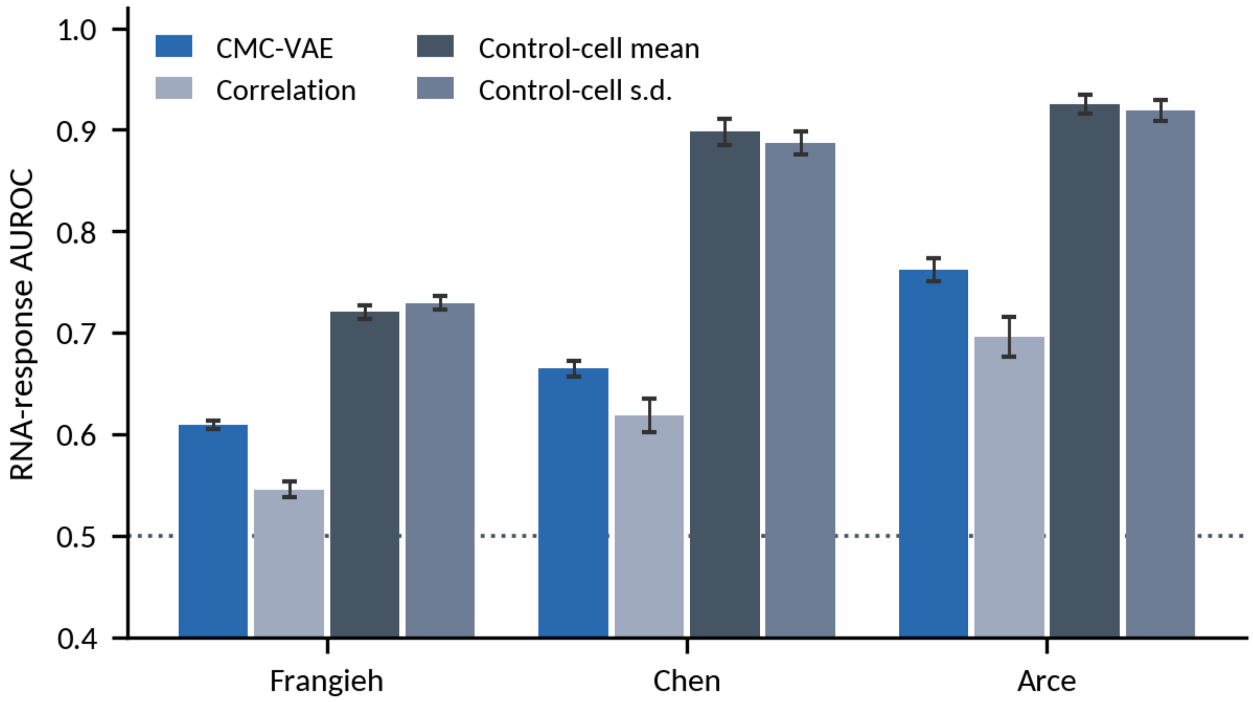


**Supplementary Fig. 9 | Knockout-invariant expression rankings.** RNA-response AUROC of CMC-VAE and the correlation baseline compared with rankings that use no perturbation information: each gene's mean or s.d. of expression in control cells. Because responsive genes share expression features across knockouts, such rankings reach high AUROC; they are references rather than perturbation-specific predictors. Error bars, s.e.m. (CMC-VAE, correlation) or bootstrap s.e. (control-cell rankings); dotted line, AUROC = 0.5.

Decoder-guided operation. The exogenous shift contributed about 0.01% of the norm of the predicted response in the development screens (about 0.1% in Papalexi); predictions with and without the shift were perfectly correlated ($r = 1.000$). Predicted responses correlated 0.52–0.93 across knockouts, compared with 0.05–0.17 for measured responses (condition-matched DE; RNA and protein). The soft do-operation lowered the predicted expression of the targeted gene in 93%, 60% and 61% of knockouts (Frangieh, Chen and Arce), compared with 98%, 100% and 100% for the signed knockdown shift and 92%, 79% and 100% in the measured responses (condition-matched evaluation set; Supplementary Fig. 12).

Fine-tuning controls (Supplementary Fig. 10 and Supplementary Table 9). Across three independently trained models, fine-tuning raised held-out RNA pearDE from 0.01 ± 0.01 to 0.20 ± 0.05, but label-shuffled targets (0.20 ± 0.04) and screen-mean targets (0.20 ± 0.06) gave the same gain, and sign-flipped targets gave −0.09 ± 0.02. Centred pearDE (0.055 versus 0.054) and retrieval (0.553 versus 0.552) did not differ between real and label-shuffled targets. Held-out protein pearDE rose from 0.14 to 0.28 with real targets, and to 0.28 and 0.34 with label-shuffled and screen-mean targets. A model-free predictor using the mean response of the training knockouts reached a held-out RNA pearDE of 0.44. On these screens, fine-tuning therefore learns mainly the response shared across knockouts.

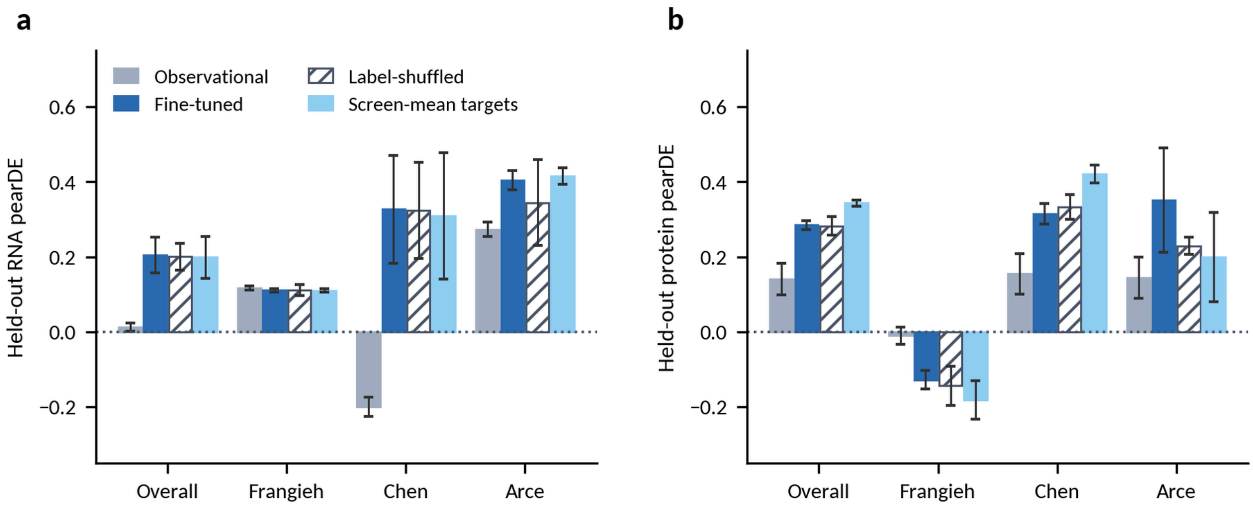


**Supplementary Fig. 10 | Fine-tuning controls on the CITE-seq screens.** Held-out RNA pearDE (a) and protein pearDE (b), pooled ('overall') and per screen, for the observational model and for fine-tuning with real, label-shuffled (hatched) and screen-mean targets. Bars, mean of three independently trained models; error bars, s.d. across models. Similar gains with label-shuffled and screen-mean targets show that, on these screens, fine-tuning mainly learns the response shared across knockouts.

Sparsity and orientation (Supplementary Table 11). In archived training runs with an L1 penalty on the latent gates and weights, the correlation between the latent graph and its transpose fell from 0.99 to as low as 0.01, and up to 87% of the strongest edge pairs were reduced to a single direction, but the retained direction agreed with CollecTRI and DoRothEA regulator-to-target orientation at chance (0.42–0.59, matching a node-level null), and RNA-response AUROC decreased. DeepSEM showed the same behaviour (49% agreement with both references). In the reported model the latent-graph sparsity term was inactive; sparsity of the projected networks arises from the decoder group-lasso. Because $G = DAD^T$, a symmetric latent adjacency yields a symmetric projected network regardless of decoder sparsity.

Penalties for direction during fine-tuning. Adding an acyclicity penalty (DAGMA; weight 0.1–100) or an asymmetry penalty to the fine-tuning objective did not orient the graph either. The projected RNA network remained nearly symmetric (correlation with its transpose ≥ 0.99, and 0.96 for the strongest asymmetry penalty with screen-mean targets), and the fraction of CollecTRI regulator–target pairs with a larger forward than reverse weight stayed at 0.51–0.55 (observational model 0.54), with the same values under label-shuffled and screen-mean targets. Held-out retrieval (0.53–0.57) and centred pearDE did not differ from those controls for any penalty (paired $P > 0.2$). Agreement with the signs of all signed CollecTRI edges was nominally higher than with label-shuffled targets for two DAGMA weights (by 0.04 and 0.05), but not higher than with screen-mean targets for any penalty.

Cognate map versus raw correlation. Raw RNA–protein correlation in the training cells gave a cognate enrichment of 8.22-fold (CMC-VAE 8.20-fold; totalVI 3.02-fold), with a median cognate rank of 3.4% (CMC-VAE 9.1%). CMC-VAE's map was more concentrated (Gini coefficient 0.74 versus 0.63; 84% versus 4% of entries below 0.1% of the maximum), but the cognate transcript was the top-ranked transcript for none of the 74 proteins (correlation, 12%).

#### Supplementary Note 16: Genome-scale Perturb-seq data

We used the essential-gene CRISPRi Perturb-seq screens in K562 cells (Replogle et al.[15]) and Jurkat cells (Nadig et al.[16]), from the preprocessed release of both screens: Jurkat (184,470 cells; 12,013 non-targeting) and K562 (188,590 cells; 10,691 non-targeting). Expression was log1p of counts normalized to $10^4$. Features were the 3,000 genes shared by both lines (1,685 perturbation targets plus 1,315 highly variable genes). Target genes were split 70/30 by gene (seed 42), applying the same split to both lines. Perturbations with at least 30 cells whose target was among the features were evaluated: 1,436 in Jurkat (1,008 training/428 held-out) and 1,310 in K562 (911/399), with a median of 262 and 192 DE genes per perturbation; the target gene itself decreased by a median of 0.42 and 0.44 log units.

Model and fine-tuning. The RNA-only CMC-VAE used the same latent structural model and decoder constraints as the CITE-seq model and was trained on the non-targeting cells and the cells of training perturbations (80/10/10 cell split for monitoring) (seeds 42, 1 and 2 for each line; models pretrained on control cells only were trained in the same way). Knockdowns were applied as a signed shift of the exogenous input, the minimum-norm change that sets the decoded expression of the target to zero. The soft do-operation (edge scaling) used for the CITE-seq screens produced predictions that were dominated by the shared response on these screens (raw pearDE 0.37, retrieval 0.49), so it was not used. For each objective, the number of steps was selected on an inner 30% split of training genes using the real-target loss, and the model was then refitted on all training genes.

Results (Supplementary Table 10). Before fine-tuning, predictions were not gene-specific (retrieval 0.52 and 0.51; centred pearDE ≈ 0), and the mean-response predictor reached raw pearDE 0.42 (Jurkat) and 0.39 (K562). Every fine-tuning objective exceeded its label-shuffled control in both retrieval and centred pearDE (paired $P \leq 3 \times 10^{-7}$), with consistent gaps across models. Gains required at least about 100 training perturbations; with 5–25 training genes all objectives remained close to chance. Fine-tuning on all perturbations of one line and evaluating the 324 genes that are held out in the other line, and perturbed in both, gave retrieval 0.56 (K562 to Jurkat) and 0.58 (Jurkat to K562) versus 0.50 and 0.52 for shuffled targets, whereas copying the measured response of the other line gave 0.74 and 0.76.

Comparison with DeepSEM and supervised predictors. DeepSEM was trained on the same cells as each CMC-VAE model, with the same features, and fine-tuned with a mask of its 73,728 largest-magnitude entries, using the same loss, objectives, label-shuffled arms, stopping rule and evaluation. Its retrieval gain over label-shuffled fine-tuning was smaller than that of CMC-VAE for both objectives in both lines (difference of gains +0.023 and +0.034 in Jurkat, +0.037 and +0.043 in K562 for the standard and contrastive objectives; $q = 0.02$, $6 \times 10^{-4}$, $2 \times 10^{-4}$ and $2 \times 10^{-4}$), and with the contrastive objective CMC-VAE also had a larger centred-pearDE gain (+0.027 in Jurkat and +0.041 in K562; $q \leq 6 \times 10^{-5}$). Ridge regression from each target's co-expression profile or its CMC-VAE decoder loadings reached retrieval of about 0.64 (centred pearDE about 0.17), exceeding latent-graph fine-tuning, as did GEARS, the linear baseline of Ahlmann-Eltze et al. and ridge regression on scGPT gene embeddings (Supplementary Fig. 11 and Supplementary Table 10).

**Robustness to the DE definition.** Redefining DE for the Perturb-seq and Papalexi screens with Benjamini–Hochberg-corrected Mann–Whitney tests ($q < 0.05$) or with pseudobulk DE (moderated t, $q < 0.05$), with the same LFC floor, reduced the median number of DE genes per perturbation from 262 and 192 to 90–129 (Jurkat and K562). The gain of fine-tuning over label-shuffled targets in held-out centred pearDE was +0.036 and +0.049 in Jurkat and +0.079 and +0.081 in K562 (BH and pseudobulk; $q \leq 10^{-4}$;

originally +0.033 and +0.050), CMC-VAE retained a larger retrieval gain than DeepSEM in every comparison, and the gain after control-only pretraining remained positive. On Papalexi, RNA-response AUROC was 0.72 and 0.70 for CMC-VAE, 0.68 for correlation, 0.58–0.59 for totalVI and 0.66–0.67 for DeepSEM; the knockout-specific end points remained not significantly different between methods (q ≥ 0.26).

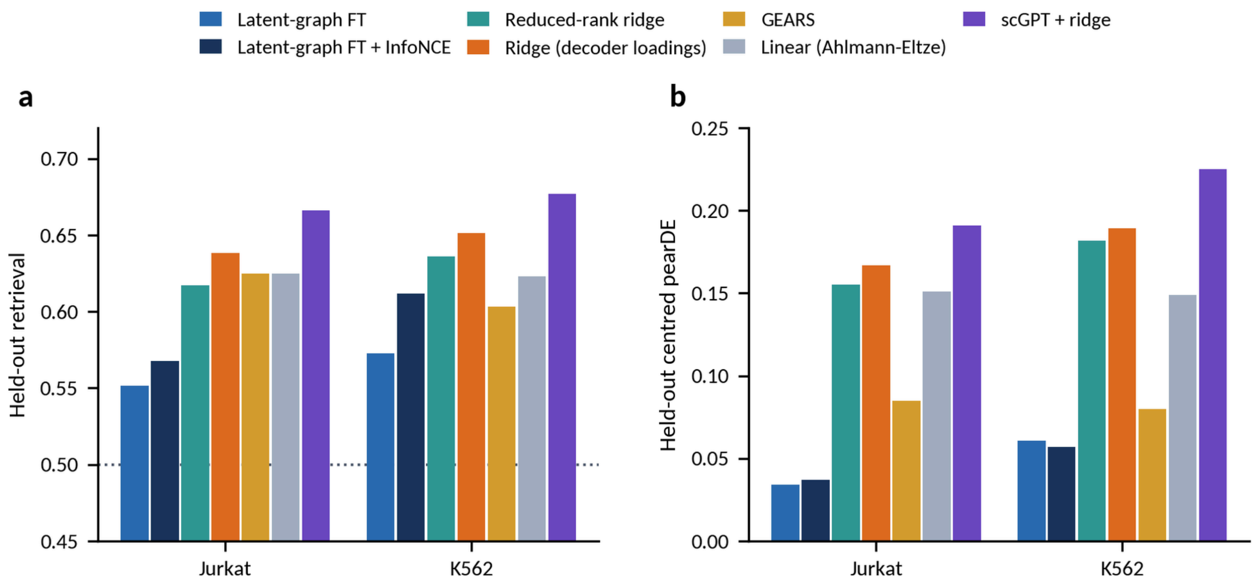


**Supplementary Fig. 11 | Latent-graph fine-tuning and supervised predictors on Perturb-seq.** Held-out retrieval (a) and mean-centred pearDE (b) on the Jurkat and K562 screens for latent-graph fine-tuning (standard and InfoNCE objectives), reduced-rank ridge regression and ridge regression from each target's CMC-VAE decoder loadings, GEARS, the linear baseline of Ahlmann-Eltze et al. and ridge regression on scGPT gene embeddings, all scored on the same held-out genes. CMC-VAE values are means of three models. Dotted line, chance.

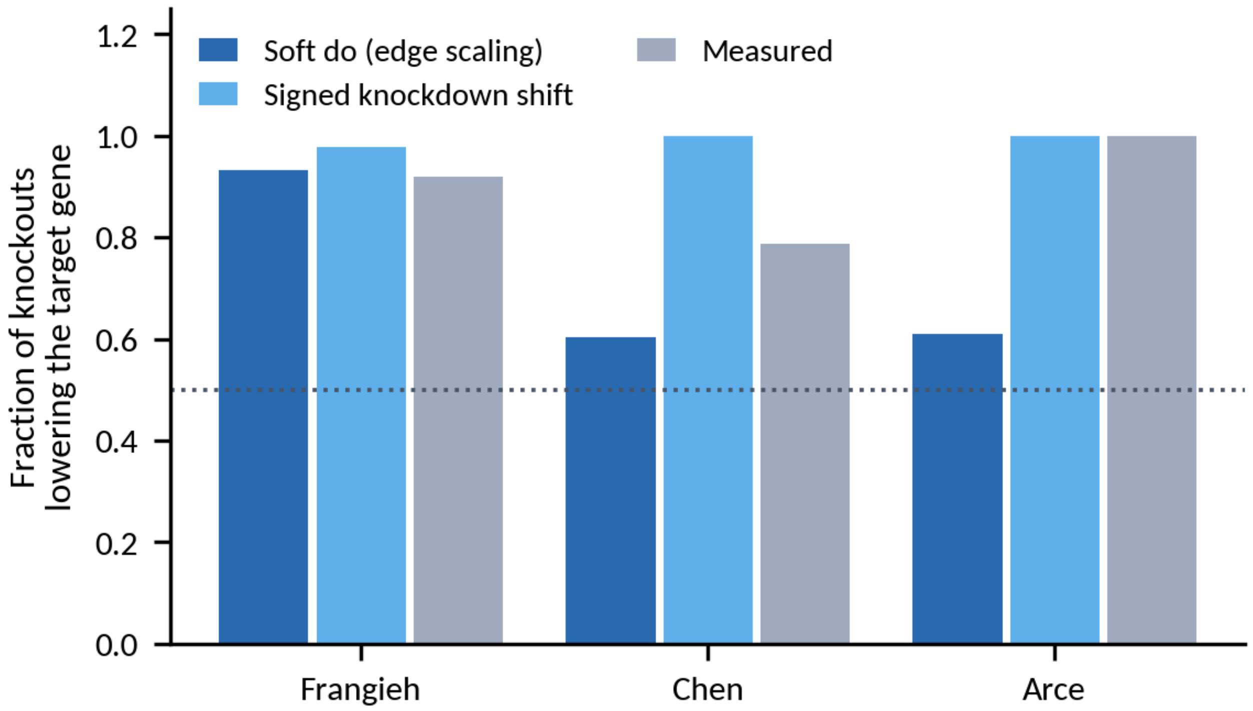


**Supplementary Fig. 12 | Sign of the in silico knockout on its own target.** Fraction of knockouts for which the predicted change of the targeted gene is negative, for the soft do-operation used for zero-shot ranking (edge scaling with absolute decoder loadings), the signed knockdown shift used for Perturb-seq fine-tuning, and the measured responses, in the three development screens (174, 122 and 23 knockouts; mean of three models). Dotted line, 0.5.

### Supplementary Tables

**Supplementary Table 1 | Training-cohort sweep.** Mean RNA-response AUROC across the three development CRISPR screens (Frangieh, Chen, Arce). "All cells" uses every quality-control-passing cell of each combination. All 15 non-empty cohort combinations are listed, followed by two additional final-pipeline entries (the Hao + Stephenson model trained with the final vocabulary pipeline, and a COMBAT-only run under that pipeline), for 17 rows in total. Sweep entries use a unified pipeline for a fair comparison across combinations.

| k | Training cohorts | All cells |
|---|---|---|
| 1 | Su | 0.669 |
| 1 | Stephenson | 0.660 |
| 1 | COMBAT | 0.654 |
| 1 | Hao | 0.612 |
| 2 | Stephenson + Su | 0.655 |
| 2 | COMBAT + Su | 0.653 |
| 2 | Stephenson + COMBAT | 0.649 |
| 2 | Hao + Stephenson | 0.635 |
| 2 | Hao + COMBAT | 0.620 |
| 2 | Hao + Su | 0.631 |
| 3 | Hao + Stephenson + Su | 0.642 |
| 3 | Stephenson + COMBAT + Su | 0.641 |
| 3 | Hao + Stephenson + COMBAT | 0.633 |

| k | Training cohorts | All cells |
|---|---|---|
| 3 | Hao + COMBAT + Su | 0.627 |
| 4 | Pooled (all four) | 0.629 |
| 2 | **Hao + Stephenson (final model)** | **0.667** |
| 1 | COMBAT only | 0.644 |

**Supplementary Table 2 | CMC-VAE and totalVI evaluated on identical held-out cells.** Protein-imputation, cognate-map and reference-network metrics are defined in Methods. Values are descriptive point estimates unless uncertainty is shown. The RNA asymmetry row describes the specified model-derived score matrices, not ground-truth causal orientation.

| Metric | totalVI | CMC-VAE |
|---|---|---|
| RNA-to-protein imputation (per-protein Pearson's *r*, CLR) | **0.502** | 0.403 |
| Cognate translation fold | 3.0 | **8.2** |
| Translation-map concentration (Gini coefficient) | 0.54 | **0.74** |
| Co-expression AUROC | **0.977** | 0.976 |
| CollecTRI AUROC | 0.574 | **0.632** |
| DoRothEA AUROC | 0.603 | **0.678** |
| RNA-to-RNA orientation asymmetry | 1.000 | ≈1.00 |

**Supplementary Table 3 | Synthetic orientation accuracy** (n = $10^4$ samples). Estimators use the information specified in Methods, including oracle latent access or decoder inversion where applicable. Values are reported point estimates. 0.5 is the random-orientation reference.

| Noise family | Our read-out ($DAD^\top$) | DirectLiNGAM on $z$ | DirectLiNGAM after decoding |
|---|---|---|---|
| Gaussian | 0.50 | 0.60 | 0.56 |
| Laplace | 0.49 | **1.00** | **1.00** |
| Student's *t*(8) | 0.49 | **1.00** | **1.00** |

**Supplementary Table 4 | Architecture and regularization variants evaluated in Frangieh.** Co-expression AUROC, cognate enrichment, RNA pearDE, forward-to-reverse asymmetry and protein AUROC are defined in Methods. The sweep baseline uses a feature-norm decoder, which differs from the final group-lasso configuration. Cognate enrichment uses a fixed strict-symbol pair set and therefore differs from the main 74-pair analysis. All 27 configurations were evaluated for the perturbation metrics (RNA pearDE and protein AUROC). For the strongest exogenous-noise bottleneck (L0 = 40), whose exogenous representation had largely collapsed (about five active exogenous dimensions of 128), the reference-network metrics (co-expression AUROC, cognate enrichment and forward-to-reverse asymmetry) were not evaluated; the dashes in that row mark these three unevaluated metrics, not missing data.

| Group | Configuration | Co-expr. | Cognate | RR pearDE | Asym. | RP AUROC |
|---|---|---|---|---|---|---|
| Base (feature-norm decoder) | Prior off (β = 2, free bits 1.0) | 0.935 | 25.4 | 0.034 | 1.02 | 0.670 |
| KL/free bits | β = 4 | 0.936 | 31.3 | 0.005 | 0.98 | 0.701 |
| | Free bits 0.0 | 0.943 | 23.1 | −0.013 | 0.99 | 0.706 |
| | Free bits 0.3 | 0.936 | 25.8 | −0.002 | 1.04 | 0.711 |
| | β = 4, free bits 0.3 | 0.939 | 24.9 | −0.003 | 1.00 | 0.692 |
| ε-bottleneck ($L_0$) | 0.5 | 0.933 | 21.7 | −0.071 | 0.97 | 0.618 |
| | 1 | 0.939 | 18.5 | −0.033 | 0.97 | 0.692 |
| | 2 | 0.911 | 13.6 | −0.088 | 0.97 | 0.566 |
| | 5 | 0.840 | 15.0 | −0.117 | 1.00 | 0.647 |

| Group | Configuration | Co-expr. | Cognate | RR pearDE | Asym. | RP AUROC |
|---|---|---|---|---|---|---|
| | 10 | 0.857 | 12.4 | 0.074 | 1.02 | 0.473 |
| | 20 | 0.789 | 7.2 | 0.007 | 1.29 | 0.524 |
| | 40 | – | – | −0.015 | – | 0.552 |
| Decoder constraint | Pathway mask (soft 0.1) | 0.963 | 24.0 | −0.003 | 1.02 | 0.596 |
| | Pathway mask (soft 0.3) | 0.960 | 41.2 | −0.046 | 1.01 | 0.673 |
| | Mask + bottleneck (0.02) | 0.937 | 19.4 | −0.070 | 0.94 | 0.638 |
| | Mask + bottleneck (0.10) | 0.933 | 20.8 | −0.021 | 0.98 | 0.602 |
| | Pathway mask (frozen) | 0.581 | 2.9 | 0.086 | 0.99 | 0.726 |
| Nonlinear SCM | tanh (initialization 0.3) | 0.901 | 31.6 | −0.060 | 0.91 | 0.572 |
| | ReLU (initialization 0.3) | 0.896 | 22.9 | 0.003 | 1.00 | 0.585 |
| | tanh (initialization 0.5) | 0.903 | 35.9 | −0.090 | 0.91 | 0.541 |
| Symmetric latent | 128/128 | 0.938 | 18.6 | −0.037 | 1.00 | 0.560 |
| Seeds | Seed 1 | 0.948 | 21.6 | −0.024 | 1.00 | 0.647 |
| | Seed 2 | 0.939 | 32.8 | 0.015 | 1.00 | 0.671 |
| Directed prior | Weight 500 | 0.939 | 12.6 | −0.039 | 1.02 | 0.716 |
| | Weight 750 | 0.749 | 13.3 | −0.079 | 7.9 | 0.708 |
| | Weight 1,000 | 0.767 | 10.2 | −0.014 | 178 | 0.613 |
| | Weight 1,500 | 0.758 | 21.7 | −0.029 | 798 | 0.551 |

**Supplementary Table 5 | Additional objectives and forward–reverse prediction differences.** Network symmetry is the correlation of the projected RNA scores with their transpose. RNA pearDE differences are reported in Frangieh, Chen and Arce order. Each row describes a particular trained variant; differences are diagnostic comparisons, not a universal test of causal identification. The reversal-loss objective is distinct from the final observational training objective.

| Lever | Mechanism | corr($G_{rr}$, $G_{rr}^{\top}$) | $A_{rr}$ forward − reverse (F/C/A) | Interpretation |
|---|---|---|---|---|
| Baseline | None (final model) | 1.000 | 0.006/−0.013/0.015 | Undirected GRN |
| iVAE: cell type | Auxiliary prior | 0.707 | −0.120/0.013/−0.050 | Asymmetric, inconsistent |
| iVAE: donor | Auxiliary prior | 0.910 | −0.088/0.136/0.316 | Asymmetric; breaks $W_{rp}$ |
| iVAE: disease | Auxiliary prior | 0.557 | 0.074/0.016/−0.033 | Asymmetric; breaks $W_{rp}$ |
| Non-Gaussian | Laplace ε prior | 0.983 | −0.012/0.021/0.036 | Symmetric, inert |
| Residual independence (TC) | TC penalty (λ = 1) | 0.998 | −0.041/0.017/0.004 | Symmetric, inert |
| Residual independence (HSIC) | HSIC penalty | 0.998 | −0.002/−0.002/−0.008 | Symmetric, inert |
| Reversal loss | Reversal loss (λ = 10) | 0.999 | 0.005/−0.001/0.023 | Symmetric, inert |

**Supplementary Table 6 | Conditional priors and fitted-operator summaries.** Rows compare priors conditioned on the joint exogenous vector with priors conditioned on RNA latents only. Columns give projected RNA symmetry, the normalized RNA-operator distance, maximum absolute RNA-to-protein weight, and RNA and protein pearDE in Frangieh, Chen and Arce order. Operator summaries are single-model values. Prediction columns use the development evaluation (cell-level DE with the uncorrected Frangieh controls; Methods). The RNA-only variants use the specified unconditional protein prior.

| Model | corr($G_{rr}$, $G_{rr}^{\top}$) | $A_{rr}$-amp. (%) | max $\lvert W_{rp}\rvert$ | RR pearDE (F/C/A) | RP pearDE (F/C/A) |
|---|---|---|---|---|---|
| CMC-VAE (final model) | 1.00 | 77 | $2.3 \times 10^{-2}$ | 0.03/−0.13/0.05 | 0.69/0.06/0.28 |
| iVAE: cell type (joint) | 0.71 | 12 | $8.6 \times 10^{-4}$ | −0.02/0.06/0.01 | 0.59/0.00/0.25 |

| Model | corr($G_{rr}$, $G_{rr}^{\top}$) | $A_{rr}$-amp. (%) | max $\lvert W_{rp} \rvert$ | RR pearDE (F/C/A) | RP pearDE (F/C/A) |
|---|---|---|---|---|---|
| iVAE: cell type (RNA only) | 0.78 | 13 | $1.0 \times 10^{-3}$ | −0.03/0.08/−0.04 | 0.54/0.00/0.35 |
| iVAE: donor (joint) | 0.91 | 34 | $1.0 \times 10^{-3}$ | 0.02/0.05/0.02 | −0.10/0.07/0.26 |
| iVAE: donor (RNA only) | 0.80 | 25 | $9.6 \times 10^{-4}$ | 0.04/0.10/0.05 | 0.12/0.09/0.21 |
| iVAE: disease (joint) | 0.56 | 12 | $7.6 \times 10^{-4}$ | 0.01/0.24/−0.14 | 0.15/−0.03/0.04 |
| iVAE: disease (RNA only) | 0.46 | 9 | $1.1 \times 10^{-3}$ | −0.04/0.25/−0.10 | 0.35/0.07/−0.30 |

**Supplementary Table 7 | Perturbation cohorts.** Evaluated perturbation units (174 / 122 / 23) are the RNA-to-RNA–evaluable set under the condition-matched pseudobulk definition (target gene in the model vocabulary and ≥3 DE genes). The number entering each metric depends on the modality, because AUROC requires ≥3 DE features and signed pearDE requires ≥5: RNA-to-RNA AUROC 174 / 122 / 23 and pearDE 174 / 105 / 19 (DE genes); RNA-to-protein AUROC 48 / 102 / 19 and pearDE 16 / 83 / 18 (DE proteins). The screening denominators in Supplementary Note 11 use the cell-level definition and are used only for the threshold analysis. The Papalexi ECCITE-seq screen (THP-1; CRISPR–Cas9 KO; 13 evaluable knockouts; 4 measured proteins) was withheld from development and used only for the final test.

| Cohort | Evaluable knockouts | Overlapping proteins | Assay | Role |
|---|---|---|---|---|
| Frangieh[17] | 174 | 7 | Perturb-CITE-seq (CRISPR–Cas9 KO) | Development (RNA and protein) |
| Chen | 122 | 78 | Perturb-icCITE-seq (CRISPR–Cas9 KO; ic, intracellular) | Development (RNA and protein) |
| Arce | 23 | 84 | Perturb-CITE-seq (CRISPRi) | Development (RNA and protein) |

**Supplementary Table 8 | Final observational model across three seeds.** Values are mean ± s.d. across the three training seeds (42, 1 and 2). Perturbation metrics use the development evaluation (cell-level DE with the uncorrected Frangieh controls) and differ from Fig. 3. The cognate translation fold of 9.8 ± 1.8 is the three-seed mean (individual seeds 8.2, 11.7 and 9.4, all on the 74-pair cognate set); the 8.2-fold value quoted in the main text is the deployed final checkpoint (seed 42) on the same 74-pair set — that is, the single-seed value whose three-seed average is reported here. The other main-text reference-network values (co-expression 0.98, STRING 0.76, DoRothEA 0.68, CollecTRI 0.63) are the corresponding final-checkpoint values and differ from these three-seed means only by rounding and seed variation.

| Metric | Mean ± s.d. | Metric | Mean ± s.d. |
|---|---|---|---|
| Co-expression AUROC | 0.981 ± 0.003 | RNA-to-RNA AUROC | 0.681 ± 0.003 |
| STRING AUROC | 0.744 ± 0.010 | RNA-to-protein AUROC | 0.535 ± 0.017 |
| DoRothEA AUROC | 0.672 ± 0.005 | RNA-to-RNA pearDE | −0.021 ± 0.020 |
| CollecTRI AUROC | 0.628 ± 0.003 | RNA-to-protein pearDE | 0.333 ± 0.026 |
| Cognate translation fold | 9.8 ± 1.8 | | – |

**Supplementary Table 9 | Fine-tuning controls on the CITE-seq screens.** Held-out values pooled over the three screens (RNA pearDE, n = 87; retrieval, n = 98; protein pearDE, n = 38), mean ± s.d. across three independently trained CMC-VAE models (seeds 42, 1 and 2). Retrieval chance is 0.5. Metrics are defined in Methods and Supplementary Note 15.

| Target | RNA pearDE | Centred pearDE | Retrieval | Protein pearDE |
|---|---|---|---|---|
| Observational (no fine-tuning) | 0.01 ± 0.01 | 0.001 ± 0.019 | 0.546 ± 0.014 | 0.14 ± 0.04 |
| Real targets | 0.20 ± 0.05 | 0.055 ± 0.026 | 0.553 ± 0.014 | 0.28 ± 0.01 |
| Label-shuffled targets | 0.20 ± 0.04 | 0.054 ± 0.021 | 0.552 ± 0.015 | 0.28 ± 0.02 |
| Screen-mean targets | 0.20 ± 0.06 | 0.045 ± 0.020 | 0.556 ± 0.015 | 0.34 ± 0.01 |
| Sign-flipped targets | −0.09 ± 0.02 | −0.007 ± 0.015 | 0.542 ± 0.012 | −0.37 ± 0.06 |

**Supplementary Table 10 | Large-scale Perturb-seq: fine-tuning, controls and baselines.** Held-out target genes (Jurkat n = 428, K562 n = 399); mean ± s.d. across three models per line (DeepSEM: one per CMC-VAE model, trained on the same cells). Label-shuffled values are means of three draws. External predictors (GEARS, linear baseline, scGPT + ridge) are single values. Retrieval chance is 0.5. See Supplementary Note 16.

| Predictor | Jurkat retrieval | Jurkat centred pearDE | Jurkat pearDE | K562 retrieval | K562 centred pearDE | K562 pearDE |
|---|---|---|---|---|---|---|
| CMC-VAE, observational | 0.524 ± 0.003 | 0.001 ± 0.006 | 0.115 ± 0.016 | 0.506 ± 0.004 | −0.004 ± 0.003 | 0.105 ± 0.010 |
| CMC-VAE, fine-tuned | 0.552 ± 0.009 | 0.034 ± 0.006 | 0.248 ± 0.004 | 0.573 ± 0.002 | 0.061 ± 0.003 | 0.233 ± 0.003 |
| CMC-VAE, label-shuffled | 0.516 ± 0.004 | 0.001 ± 0.004 | 0.231 ± 0.002 | 0.521 ± 0.001 | 0.011 ± 0.001 | 0.200 ± 0.002 |
| CMC-VAE, fine-tuned (InfoNCE) | 0.568 ± 0.008 | 0.037 ± 0.005 | 0.187 ± 0.050 | 0.612 ± 0.014 | 0.057 ± 0.009 | 0.110 ± 0.015 |
| CMC-VAE, InfoNCE label-shuffled | 0.501 ± 0.020 | 0.002 ± 0.005 | 0.151 ± 0.062 | 0.518 ± 0.004 | 0.004 ± 0.003 | 0.064 ± 0.018 |
| CMC-VAE (control-only pretraining), fine-tuned | 0.557 ± 0.008 | 0.022 ± 0.004 | 0.175 ± 0.015 | 0.542 ± 0.007 | 0.019 ± 0.004 | 0.158 ± 0.011 |
| CMC-VAE (control-only pretraining), label-shuffled | 0.530 ± 0.006 | 0.002 ± 0.003 | 0.157 ± 0.011 | 0.505 ± 0.006 | −0.009 ± 0.003 | 0.137 ± 0.012 |
| DeepSEM, observational | 0.514 ± 0.004 | −0.002 ± 0.002 | 0.015 ± 0.021 | 0.523 ± 0.003 | −0.026 ± 0.002 | 0.051 ± 0.013 |
| DeepSEM, fine-tuned | 0.542 ± 0.005 | 0.005 ± 0.004 | 0.250 ± 0.056 | 0.549 ± 0.005 | −0.032 ± 0.004 | 0.253 ± 0.023 |
| DeepSEM, label-shuffled | 0.529 ± 0.003 | −0.019 ± 0.008 | 0.262 ± 0.019 | 0.534 ± 0.002 | −0.054 ± 0.002 | 0.269 ± 0.003 |
| DeepSEM, fine-tuned (InfoNCE) | 0.521 ± 0.006 | 0.002 ± 0.005 | 0.034 ± 0.027 | 0.552 ± 0.002 | 0.008 ± 0.006 | 0.043 ± 0.013 |
| DeepSEM, InfoNCE label-shuffled | 0.487 ± 0.006 | −0.007 ± 0.007 | 0.004 ± 0.030 | 0.501 ± 0.002 | −0.003 ± 0.006 | 0.016 ± 0.013 |
| Ridge, decoder loadings | 0.638 ± 0.004 | 0.167 ± 0.007 | 0.442 ± 0.002 | 0.651 ± 0.002 | 0.189 ± 0.002 | 0.432 ± 0.001 |
| Ridge, co-expression profile | 0.636 ± 0.000 | 0.161 ± 0.000 | 0.436 ± 0.000 | 0.639 ± 0.000 | 0.163 ± 0.000 | 0.419 ± 0.000 |
| Reduced-rank ridge (decoder span) | 0.617 ± 0.006 | 0.155 ± 0.007 | 0.419 ± 0.007 | 0.636 ± 0.005 | 0.182 ± 0.005 | 0.411 ± 0.002 |
| 10-NN, decoder loadings | 0.603 ± 0.012 | 0.129 ± 0.011 | 0.380 ± 0.009 | 0.617 ± 0.008 | 0.148 ± 0.008 | 0.362 ± 0.005 |
| GEARS | 0.625 | 0.085 | — | 0.603 | 0.080 | — |

| | | | | | | |
|---|---|---|---|---|---|---|
| Linear baseline (Ahlmann-Eltze et al.) | 0.625 | 0.151 | — | 0.623 | 0.149 | — |
| scGPT embeddings + ridge | 0.666 | 0.191 | — | 0.677 | 0.225 | — |

**Supplementary Table 11 | Latent-graph sparsity creates asymmetry, not orientation.** Archived training runs with an L1 penalty on latent gates and weights (sum reduction) compared with the reported model and DeepSEM. Orientation is the fraction of reference TF⟶target pairs with |G[tf, target]| > |G[target, tf]| among pairs with non-negligible weight (> $10^{-6}$ × maximum); node-level null, TF⟶random non-target pairs. F/C/A, Frangieh/Chen/Arce. RNA AUROC uses the development (cell-level) evaluation and therefore differs from Fig. 3.

| Model | corr(A, $A^T$) | Strong pairs one-sided | CollecTRI / DoRothEA fwd > rev | RNA AUROC (F/C/A) |
|---|---|---|---|---|
| CMC-VAE, reported (no latent L1) | 0.994 | 0% | 0.54 / 0.50 | 0.63/0.67/0.74 |
| Latent L1, λ = 0.005 | 0.80 | 21% | 0.51 / 0.53 | 0.58/0.65/0.69 |
| Latent L1, λ = 0.05 | 0.94 | 27% | 0.51 / 0.50 | 0.57/0.64/0.69 |
| Latent L1, λ = 0.2 | 0.01 | 87% | 0.55 / 0.59 (null 0.57 / 0.59) | 0.58/0.64/0.68 |
| Latent-only L1, λ = 0.005–0.2 | 0.42 to 0.02 | 9–79% | 0.42–0.56 / 0.45–0.52 | 0.55/0.60/0.66 |
| Combined strong sparsity | 0.01 | 16% | 0.49 / 0.48 | 0.50/0.53/0.59 |
| DeepSEM (L1 α = 100) | 0.32 | 50% | 0.49 / 0.49 | 0.57/0.68/0.76 |
